# Telomeric amplicons of *SUL1* and Y’ in yeast are generated by microhomology-mediated break induced replication occurring *in cis*

**DOI:** 10.64898/2026.04.07.716220

**Authors:** Bonita J. Brewer, Rebecca Martin, Elizabeth Ramage, Celia Payen, Sara C. Di Rienzi, Yang Zhao, Kelsey Zane, Jocelyn Verhey, Miranda Zalusky, Danny E. Miller, Giang T. Ong, Jamie L. McKee, Gina M. Alvino, Maitreya J. Dunham, M. K. Raghuraman

**Affiliations:** Department of Genome Sciences, University of Washington, Seattle, Washington; Division of Genetic Medicine, Department of Pediatrics, University of Washington and Seattle Children’s Hospital, Seattle, Washington; Department of Laboratory Medicine and Pathology, University of Washington, Seattle, Washington; Brotman Baty Institute for Precision Medicine, University of Washington, Seattle, Washington

## Abstract

Gene amplification is a potent driver of evolution and is thought to contribute to genetic diseases, including cancer. The yeast *Saccharomyces cerevisiae* is a powerful organism for understanding amplification mechanisms. When yeast is grown long term in sulfate-limiting chemostats, amplification of the gene that encodes the primary sulfate transporter, *SUL1*, is a common outcome. Here we describe a form of *SUL1* amplification in which multiple copies of the right terminal region of chromosome II are appended in tandem to a native telomere. We find this form of amplicon when we delete the origin of replication next to *SUL1* or delete a variety of genes involved in DNA metabolism. It is the only form of amplification found in a *yku70Δ* mutant suggesting that unprotected telomeres are involved. We propose that these terminal addition events occur when the unprotected 3’ G_1-3_T telomeric sequence invades a short (∼7 bp) internal telomere sequence (ITS) to begin a form of microhomology-mediated break-induced replication (mmBIR) that has been documented in type-I survivors of telomerase mutants. In addition to amplification of the right end of chromosome II we also find that telomeres containing the sub-telomeric repeat Y’ experience similar tandem amplification events and show that their formation is reduced in a *pol32Δ* mutant, a gene required for mmBIR. Within individual amplicons the ITSs and Y’s are nearly identical, suggesting that the multiple copies of the amplified region are generated in a single mmBIR event that we describe as pseudo-rolling circle mmBIR. A similar amplification event at the P-telomere of human chromosome 18 has four copies of a ∼54 kb region separated by ITSs of nearly identical size. This finding suggests that these additional copies of the terminal fragment of human chromosome 18 arose by the same pseudo-rolling circle mechanism, perhaps during a period of telomeric stress.

**AUTHOR SUMMARY:** The human genome is peppered with duplicates (or higher numbers) of segments that are located at sites both nearby and distant from the original, ancestral segments. These Copy Number Variants, or CNVs, appear to be highly variable among different individuals and are being examined with great interest as potential loci associated with genetic disease. Experimentally determining how these CNVs arise and become distributed across the genome is nearly impossible using humans. We are using budding yeast as the model organism to explore mechanisms of gene amplification. In this work we show that by destabilizing the ends of yeast chromosomes (telomeres) or by interfering with genes involved in the replication, repair, or recombination of DNA results in a specific form of segmental copy number increase that is initiated at telomeres. We propose that a telomere invades an internal chromosome site and sets up a pseudo-circular template for conservative DNA replication. The outcome is a chromosome with multiple, identical copies of a chromosome end arranged in tandem. We believe that it is also a major mechanism used by cells to repair telomeres that have become eroded during aging.

## INTRODUCTION

Gene amplification is a common adaptive outcome when cells are exposed to environmental stressors. The yeast *Saccharomyces cerevisiae* is an ideal organism to identify and study mechanisms of gene amplification because of its relatively fast generation time and the number of molecular and genetic tools available for identifying changes at the genomic level. When a chemostat is employed to limit the growth of a population of cells by controlling access to an essential nutrient, discrete samples collected over time provide reliable and reproducible molecular data for dissecting the process of gene amplification.

Gresham et al. [1] showed that growing yeast in sulfate limiting chemostats invariably selects for amplification of the region of chromosome II that contains *SUL1*, the gene that encodes the primary sulfate transporter [1, 2]. Moreover, in wild type haploid laboratory yeast the most common form of amplicon is an interstitial inverted triplication that is variable in length, but always includes the adjacent origin of replication, *ARS228* and has junctions that map to short (∼6 bp), preexisting inverted repeats that are spaced ∼80 bp apart [2]. We proposed the ODIRA (Origin Dependent Inverted Repeat Amplification) model that can explain these inverted amplicons [3]. Among the key features of ODIRA is a replication error in the form of a template switching event between the leading and lagging strands at the diverging replication forks that arose from *ARS228*. In four of the seven chemostats seeded with a yeast strain that had a deletion of this origin, the inverted amplicons were larger and included one of the adjacent origins of replication. However, in the other three chemostats, the resulting amplicon structure suggested that a different mechanism of amplification, distinguishable from ODIRA, was responsible [4].

In the current work we further explored non-ODIRA events and identified a mechanism to explain the chromosome structures that we recovered. We used strains with mutations in genes whose products are involved in DNA replication, recombination and/or maintenance and catalogued the outcomes. We found that the gene mutation with the most dramatic and reproducible shift toward a non-ODIRA mode of amplification was the deletion of *YKU70*. From chemostats of a *yku70Δ* mutant, we only recovered amplicons that extended through the telomere. Long read sequencing revealed tandem *SUL1* fragments appended directly to a telomere that was usually, but not exclusively, the right telomere of chromosome II. The junctions occurred through interactions between the telomere and centromere-proximal, short (∼7 bp) internal telomere sequences (ITSs). In addition to these *SUL1* terminal amplicons, we found extensive variations in karyotypes in the *yku70Δ* chemostats which are explained by amplification of subtelomeric elements.

While all yeast telomeres end in a G_1-3_T tract with a 3’ overhang of the G-rich strand [5, 6], their subtelomeric regions are comprised of several sequence modules, creating enough variety to distinguish most of the 32 telomeres uniquely. Just internal to the G_1-3_T tract is an X-element that is thought to serve as an origin of replication, followed by X-element combinatorial repeats [7]. Roughly half of the 32 chromosome ends have this simple structure. The other chromosome ends contain a subtelomeric Y’ element [8] inserted between the X-element repeats and the G_1-3_T tract with an additional centromere-proximal ITS of variable length. Y’-elements are classified as “long” or “short” based on whether they encode a potentially functional helicase; short Y’s have an internal deletion in this ORF. Single Y’s are found at fifteen chromosome ends. Chromosome XII in the reference genome is the exception in that each end has two Y’s in tandem—two long Y’s on the left arm and two short Y’s on the right arm.

Separating these two copies of Y’s are ITSs of variable sizes (∼10 to 250 bp).

The structural changes identified in the *yku70Δ* chemostats are similar to those found among rare survivors of telomerase-negative mutants [9, 10]. The proposed mechanism, known as Type-1 ALT (for Alternate Lengthening of Telomeres), is a form of break induced replication (BIR) that depends on Pol32, Rad51 and Rad52 and on microhomology between the telomere and internal telomeric sequences (reviewed in [11, 12]). This form of mmBIR (microhomology mediated break induced replication) stabilizes the chromosome ends by expanding the number of the Y’ elements. Yet while each of our *SUL1* and Y’ tandem amplicons recovered from the *yku70Δ* chemostat is unique, the adjacent Y’ and ITS repeats are identical and have identical junctions—a novel feature that to our knowledge has not been reported previously in the literature on telomerase mutants.

Here we detail a specific form of mmBIR where the unprotected telomere initiates conservative replication by invading an ITS. Instead of occurring in *trans*—between sisters, for example—we propose that the invasion occurs in *cis*, producing a pseudo-rolling circle form of replication generating multiple, tandem copies of the same telomeric segment (*SUL1* or Y’) from a single initiating invasion. Y’ elements are limited to yeast; however, we believe that this amplification mechanism has been conserved from yeast to humans. In a search of the human Telomere-to Telomere (T-2-T) genome sequence (genome.ucsc.edu; T2T CHM13v2.0/hs1) we identified a tandem triplication of a 54 kb telomere-adjacent region of Chr18p in which the repeats were separated by ITSs of similar size—features consistent with those of the *SUL1* and Y’ amplicons that arise in the *yku70Δ* and *ars228Δ* yeast mutants.

## RESULTS and DISCUSSION

### Gene amplification during chemostat growth of yeast in sulfate-limiting medium

A chemostat is a culture vessel with ports for the continuous, controlled addition of media (and air) and an overflow valve to collect spent medium and a representative sample of the cells from the chemostat. By limiting a single nutrient, cells can be grown continuously under stressful conditions for hundreds of generations [13]. The rate of medium inflow sets the growth rate of the culture, but individual cells that have acquired a beneficial mutation, specific to the imposed selection, will eventually sweep the population as it gives their descendants a survival advantage [1]. By sampling cells from the overflow valve, we can follow the progression of different amplification events over time.

Among the many sulfur-limiting chemostats we have run with a prototrophic haploid yeast strain (FY4), we invariably recover cells that have amplified the sulfur transporter gene *SUL1* [1, 2, 4, 14, 15]. An example of these analyses is shown in Fig 1A-D. We detect changes in karyotype by CHEF gel electrophoresis, a form of electrophoresis that separates intact yeast chromosomes. In the ethidium bromide-stained gel (Fig 1A), there is an obvious change in the size of chromosome II on day 13, followed by its slow disappearance over the course of the remaining days. Hybridizing the Southern blot with a *CEN2* probe (Fig 1B) confirms the identity of the changed chromosome and explains its apparent disappearance—with time, a second, larger version of chromosome II that co-migrates with chromosomes XIII and XVI outcompetes cells with the initial chromosome II variant. By probing the same blot with *SUL1* (Fig 1C) and comparing the hybridization intensity of *SUL1* relative to *CEN2* on the same chromosome, we estimate that the number of copies of *SUL1* present on Chr II progress from 1 to 3 to 5, without passing through copy numbers of 2 and 4.

**Figure 1:**
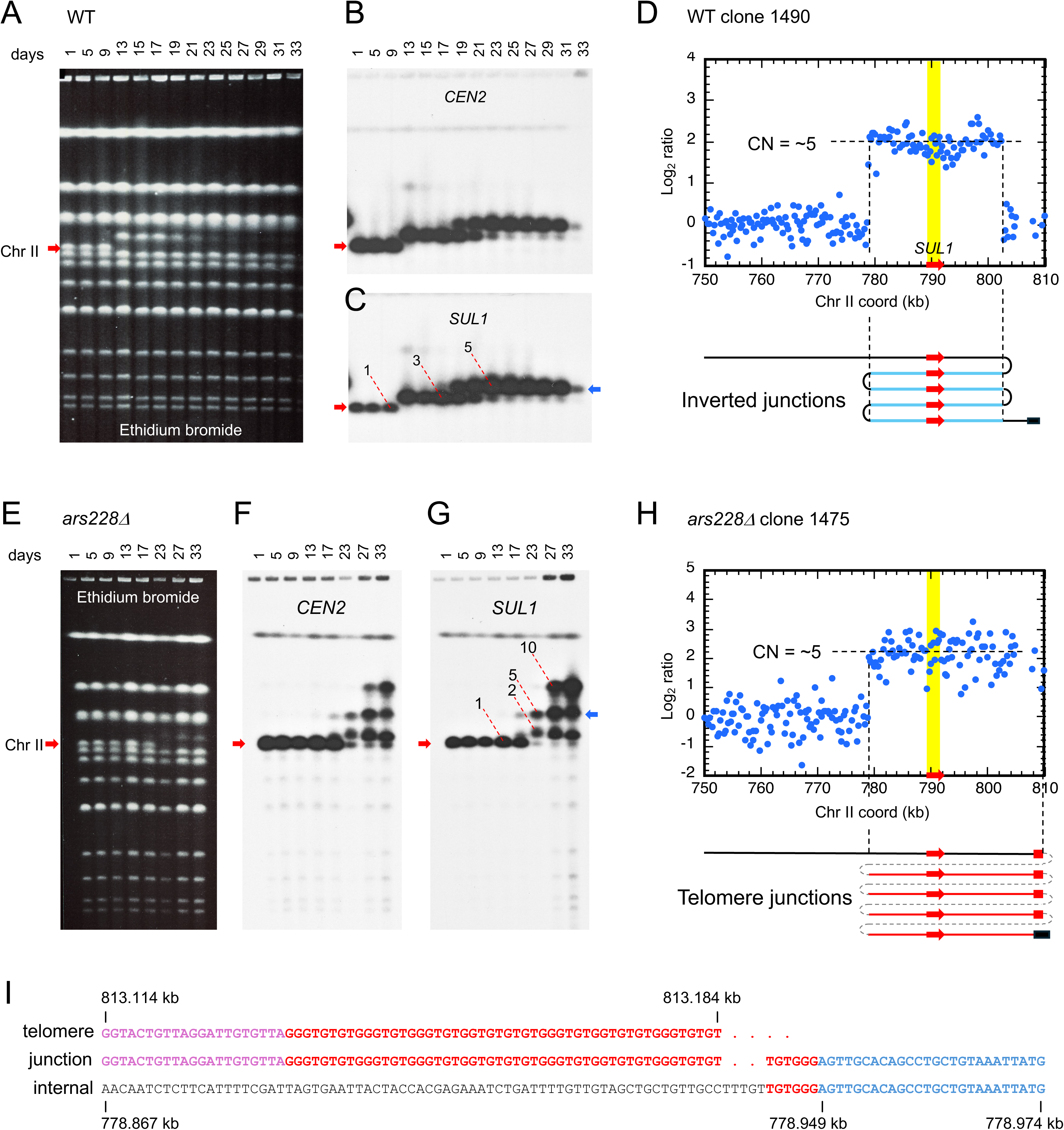
Detection of gene amplification during sulfate-limited chemostat growth. To follow karyotypic changes during ∼200 generations (33-34 days) of growth in chemostats limiting for sulfate we ran Contour-clamped Homogenous Electric Field (CHEF) gels and performed array Comparative Genome Hybridization (aCGH). Shown are examples of changes occurring in wild type (A-D) and *ars228Δ* (E-H) strains. A) Ethidium bromide-stained CHEF gel of the wild type population sampled at intervals across 34 days of growth. The expected position of chromosome II is indicated by the red arrow. Hybridizations of the CHEF gel with probes for *CEN2* (B) and *SUL1* (C) confirm step-wise changes in the size of chromosome II. The comparison of hybridization intensity of *SUL1* relative to *CEN2* indicates discrete increases in the number of copies of *SUL1* from 1 to 3 to 5 over time. D) ArrayCGH of a clone from day 34 reveals the region of chromosome II that is included in the amplification event (∼779 to 802 kb) and the number of copies (∼5; size of the terminal amplicon, blue arrow in panel C). This pattern of amplification is consistent with tandem repetition of the amplicon that alternates in orientation, leaving the distal chromosomal sequences intact. E) The ethidium bromide-stained CHEF gel of the *ars228Δ* mutant appears similar to that of the wild type strain, but hybridizations with the *CEN2* (F) and *SUL1* (G) probes indicate the persistence of multiple forms of chr II in the population. H) ArrayCGH of a clone derived from day 34 shows amplification of *SUL1* fragment that extends through the telomere and is present in five copies (blue arrow in G). I) Nanopore sequencing of clone 1475 reveals a single junction that joins the amplified fragment to the right telomere of chr II. Individual long reads confirm multiple tandem repeats.

To determine the extent of amplified flanking sequences we performed array comparative genome hybridization (aCGH, Fig 1D). In this example, we isolated a single clone (1490) from the homogeneous final population and compared genome-wide copy number of the clone to the same strain on day 0. The amplified sequences include a ∼25 kb fragment that contains the *SUL1* locus that is present at ∼5 copies (Fig 1D). These additional ∼100 kb are consistent with the size of chromosome II measured on the last day in the CHEF gel. These five copies form a tandem array of repeating units that alternate in their orientation and create junctions of head-to-head and tail-to-tail configurations similar to those found by Araya et al. [16]. In a recent study of 31 independent sulfate-limited chemostats [2] we confirmed that inverted amplicons are the predominant form of *SUL1* amplification in wild type haploid laboratory yeast strains. Among the short-read sequences from the 31 populations analyzed on the final day (day 33-35), we observed 50 unique junctions mapping between *CEN2* and *SUL1* that were consistent with a genomic inversion at short, closely spaced inverted repeats and only a small number of other types of junctions.

As a test of our proposed model for how these inverted amplicons occur (ODIRA, [3]) we had previously deleted the origin of replication at the 3’ end of *SUL1* (*ARS228*) and grown the *ars228Δ* strain in seven independent sulfate-limited chemostats [4]. Four of the seven cultures showed *SUL1* amplification patterns consistent with ODIRA, but there were three exceptions that we had not characterized in detail. In one of those three *ars228Δ* chemostats we observed chromosome II variation (Fig 1E-G) with several copy number versions (2, 5, and 10 copies) persisting through the last day. The single clone (1475) from day 33 of the *ars228Δ* chemostat that we examined by aCGH contains a similarly sized amplicon to the wild-type clone (1490), but in this case, the amplicon extends through the telomere (Fig 1H). To validate this presumed amplicon structure and to identify the precise location of the amplicon boundary, we performed long read (Oxford Nanopore) sequencing of this clone’s genome. We found a single split-read at the site that corresponds with the jump in copy number in the aCGH profile. It is a junction between the telomere of chromosome II and a unique sequence centromere-proximal to *SUL1* (Fig 1I) and is the same type of junction we found among four of the short-read sequences from the 31 chemostats of wild type haploids in which ODIRA events had predominated (S1 Fig; [2]). Several observations are consistent with the terminal, tandem repeat structure for this clone proposed in Fig 1I: the only chromosome with an altered size on the CHEF gel was chr II; sequencing returned only a single junction sequence; and individual long reads spanned multiple, tandem, direct repeats of the *SUL1* region. Therefore, deleting the origin adjacent to S*UL1* alters the response to limited sulfate and results in changes in the subtelomeric region that cannot be explained by ODIRA.

Because the non-ODIRA amplicons had junctions between a chromosomal telomere and internal telomeric sequences, we asked if a mutation affecting telomere maintenance would alter the cellular response to growth in sulfate limited conditions. Among the many proteins that bind yeast telomeric sequences is the Ku70/80 complex, which is important for telomere maintenance and genome stability [17]. Ku70/80 binds directly to chromosome ends, protecting them from degradation by exonucleases and from end-to-end fusion events. Ku70/80 also binds to telomeric DNA through its interactions with Rap1, Sir4 and the RNA component of telomerase (Tlc1) and influences telomere maintenance [18, 19]. Cells lacking either component of the Ku complex have shorter telomeric G_1-3_T tracts and show increased levels of genome instability [20–23].

After long-term growth of a *yku70Δ* strain in two sulfate-limited chemostats, we found *SUL1* amplification both cultures (*yku70Δ*-A and-B; Fig 2), which were noteworthy in that the amplicons not only extended through the right telomere of chromosome II but together they represented a minimum of six different junctions. In chemostats with wild type cells it is rare to find more than two unique junctions [2]. The multiple amplicons in chemostat *yku70Δ*-A appeared to arise concurrently each with five or more copies of the *SUL1* region, and one amplicon appeared to be located on a chromosome other than Chr II (Fig 2A,B). Because chromosome II is the primary chromosome to show a significant change in size on the CHEF gel, we concluded that the majority of these different chromosome II fragments must be appended to or incorporated into chromosome II (Fig 2C). One explanation for the arrangement of the multiple amplicons is that each of the fragments is found in a different subset of cells in the population. A second possibility is that there is a single event that contains a mixture of different-sized repeats. The results from chemostat *yku70Δ*-B (Fig 2D-F) were similar to those of *yku70Δ*-A.

**Figure 2:**
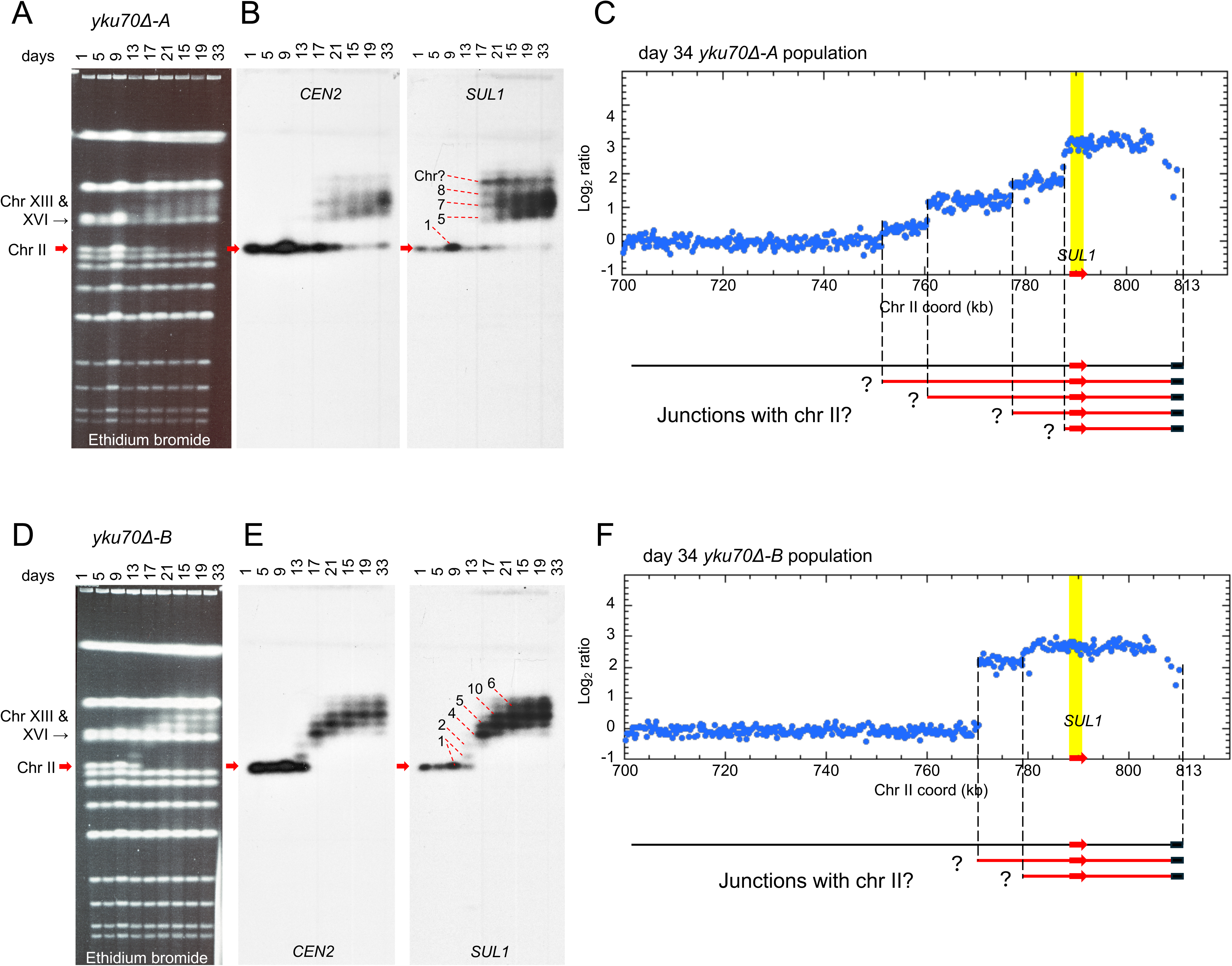
**Deletion of *yku70Δ* results exclusively in terminal amplification of *SUL1*** Chemostats *yku70Δ*-A and *yku70Δ*-B contained independent evolutions of the *yku70Δ* mutant. The ethidium bromide-stained CHEF gels (A and D) and their Southern analyses (B and E) suggest that multiple amplicons with variable numbers of copies of *SUL1* (numbers with dashed red lines) had persisted through the last day of chemostat growth. (The identity of “Chr?” is likely either chr XIII or chr XVI since that doublet decreases in intensity in the second half of the time course.) aCGH of the populations (C and F) confirm the existence of multiple unique amplicon junctions—a minimum of four in *yku70ι1*-A and two in *yku70ι1*-B. From these analyses it is unclear whether the culture is a mixture of multiple unique events, or whether the different amplicons are contiguous on the same DNA molecules.

To disentangle the different amplicon sizes found in the *yku70Δ* chemostats, and to see if they were changing over time, we isolated clones from day 17 and day 33 from *yku70Δ*-A (Fig 3A) and days 13, 17 and 33 from *yku70Δ*-B (S2A Fig). We found 33 clones with unique karyotypes, and all but one had additional copies of *SUL1*. By comparing the ethidium-stained gel with the hybridization results for *CEN2* (Fig 3B; S2B Fig) and *SUL1* (Fig 3C; S3B Fig) we observed that 1) all clones had only a single copy of chr II with some clones retaining the parental-sized chr II; 2) clones with the parental-sized chr II had amplified versions of *SUL1* on some other chromosome; and 3) many chromosomes had increased in length but the increase was not due to *SUL1* amplification. We conclude that the multiple amplicons seen in Fig 2 reflect a mixed population of cells, with different subsets having different amplicons. The explanation for the wide-spread changes in chromosome size was revealed by hybridization with a Y’ probe (Fig 3D, S2D Fig): many chromosomes had amplified copies of subtelomeric Y’ sequences. All unique bands on the CHEF gel can be accounted for by either *SUL1* or Y’ amplification or both. That these changes are selected for by growth in the sulfate-limited chemostat is illustrated by parallel analysis of independent clones from the day 0 sample (Fig 3E-G). Among these 18 clones there is only a single chromosome with a modest increase in Y’ hybridization (Fig 3G; red star).

**Figure 3:**
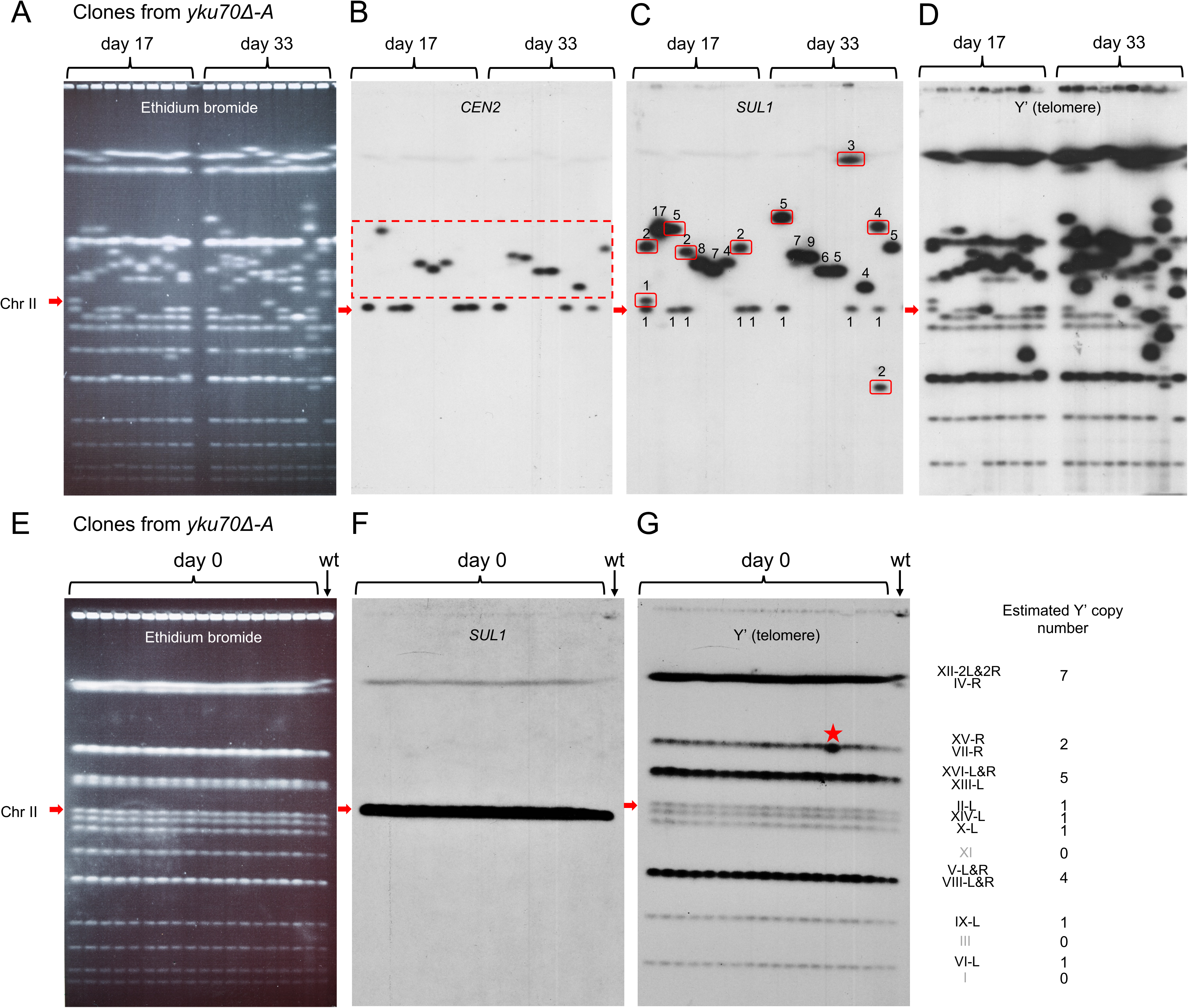
Clones from the *yku70ι1*-A chemostat have highly variable karyotypes. To determine the structure of the four terminal amplicons in the *yku70ι1*-A chemostat and *ykuΔ*-B (S2 Fig) we isolated clones from different days and analyzed them by CHEF gels and Southern hybridization. The red arrow indicates the expected position of chr II. Multiple chromosomal changes are apparent in the ethidium bromide-stained gel (A) with no two clones appearing to have the same karyotype. This conclusion is confirmed by hybridizations: B) *CEN2* hybridization reveals that many, but not all, of the clones have an altered size of chr II (red dashed box); C) *SUL1* hybridization indicates that every clone but one has an increased copy number (indicated by numbers above or below each band) of *SUL1*; however, in many cases copies of *SUL1* have moved to other chromosomes (bands highlighted by red boxes); D) Y’ hybridization reveals multiple examples of Y’ amplification that are distributed across many yeast chromosomes. The combination of *SUL1* and Y’ amplification can account for all of the karyotypic changes found in the ethidium bromide images of the *yku70ι1* CHEF gels. E and F) Eighteen clones from day 0 of *yku70ι1*-A were isolated and compared to the wild type strain using CHEF gel electrophoresis (E) and Southern blotting with the *SUL1* probe (F). The karyotypes of the mutants were indistinguishable by ethidium bromide staining. (G) Hybridization with the Y’ probe revealed a single example of Y’ amplification on chromosome XV or VII (red star) with all other chromosomes in this and other clones having a distribution of Y’ elements that were similar to estimates from the Nanopore sequencing of FY4 and the *yku70Δ* parent strain.

### Long read sequencing confirms *SUL1* amplicon structures in the yeast *yku70Δ* mutant

To identify the precise sequence of the amplicon junctions generated in the *yku70Δ* mutant strain, we performed Oxford Nanopore sequencing on the population sample from day 34 of *yku70Δ*-A. We found split reads (Fig 4A) at the same coordinates that flanked the discontinuities in copy number in the aCGH data (Fig 2C) along with four additional low-frequency junctions on chr II-R (Fig 4B). At seven of these genomic locations in the reference genome (Saccharomyces Genome Database, SGD) are short stretches of telomeric repeats (G_1-3_T/C_1-3_A) and at the eighth junction is a sequence homologous to a region just 45 bp in from the right telomere of chrII (Fig 4B). From the long reads we determined the structure of the amplicons associated with each junction: most long reads revealed tandem repeats of the telomeric *SUL1* fragment appended directly to the right telomere of chr II (Fig 4C,D). Lower frequency events involved the addition of a Y’ to the *SUL1* amplicon (Fig 4E) or translocation of the terminal *SUL1* fragment to a different chromosomal telomere (Fig 4F). For the sequences summarized in Fig 4D, the preexisting ITSs had undergone an expansion from just a few base pairs to up to nearly 220 bp (Fig 4B) and the first sixteen base pairs of the ITS was identical to the first sixteen base pairs of the chrII-R telomere (5’-GGGTGTGTGGGTGTGG-3’). This sequence identity suggests that the native chrII-R telomere was instrumental in creating the expanded G_1-3_T sequence at the *SUL1* amplicon junctions.

**Figure 4:**
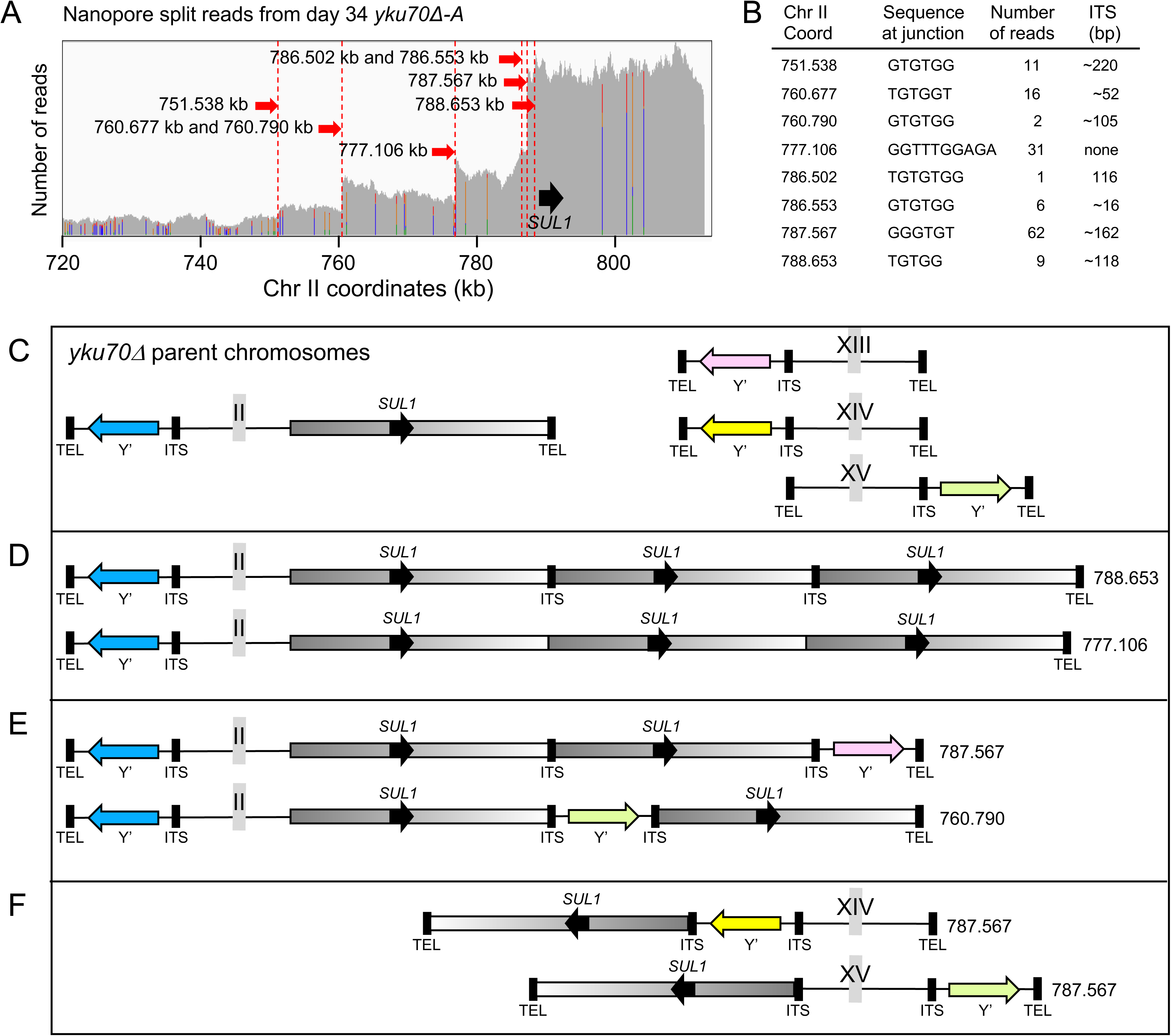
**Nanopore sequencing of the *yku70****Δ***-A chemostat population** A) A read-depth plot of the split reads from Oxford Nanopore sequencing of the *yku70Δ*-A day 34 population was consistent with the positions of copy number change that were identified in the aCGH profile (Fig 2C). B) We identified eight unique split reads from this region of chr II and found that seven of the eight junctions occurred between telomeric G_1-3_T sequences and an internal sequence centromere proximal of *SUL1*. At each internal site was a short region of G_1-3_T homology (an internal telomeric sequence, ITS). The eighth junction occurred between an internal sequence that had homology to a sequence 45 bp into the subterminal region of chr II-right. The reference genome coordinate of the junction sequence, the number of supporting reads and the length of the expanded ITS are given. C) Illustration of the telomere structure of four of the chromosomes in the parent *yku70Δ* parent strain. The colors of the Y’s are from the genome-wide analysis shown in S3A Fig. Long reads across the eight junctions provided information on the identity of the chromosome that had acquired the *SUL1* telomeric fragment, the number of repeats added, and the chr II coordinate of the amplicon junction (numbers in kb to the right of each cartoon). They fell into three classes: D) terminal addition of the *SUL1* fragment directly to the telomere or sub-telomere of chromosome II-R; E) terminal addition of the *SUL1* fragment with interspersed Y’s from other chromosome locations; and F) translocations of the terminal *SUL1* fragment to the telomeres of other chromosomes. The expanded ITSs in D and E began with 5’-GGGTGTGTGGGTGTGG-3’, the same sequence found at the junction between ChrII’s right telomere and the adjacent subtelomeric DNA.

### Structure of Y’ amplicons in the *yku70Δ* mutant

Since all clones from days 17 and 33 of the *yku70Δ*-A evolution had one or more chromosomes with additional copies of Y’ (Fig 3D), we searched Nanopore split reads near telomeres and found many instances of tandem arrays of Y’s. We wanted to address three questions: 1) is there a specific Y’ that is amplified; 2) within an array, how homogeneous are tandem Y’s and ITSs; and 3) how are amplification of Y’ and *SUL1* related? To classify Y’s and assign them accurately to chromosomes, we defined a section of the Y’ region (bounded by specific sequence tags common to all Y’s curated in SGD; “trimmed Y’s”) and determined alignments and phylogenetic relationships of the Y’s in the reference genome (Saccer3) with those in FY4 and the *yku70Δ* parent (S3 Fig). These comparisons defined 13 Y’ classes that we could assign to specific chromosomes (S3 Fig). We then used the set of trimmed Y’s from FY4 to compare individual Y’s from tandem amplicons using multiple alignments and phylogenetic trees.

Examples of four Nanopore reads that contain tandem Y’ repeats are diagrammed in Fig 5. The first three reads (Fig 5A,B,C) are composed of homogeneous Y’s that match the Y’ that is resident on that specific chromosome. The fourth read (Fig 5D) is a repeating trimer of two distinct Y’s. Because this read (as well as read 3) did not contain unique sequences, we could not definitively map it to a particular chromosome. However, based on the phylogenetic trees (S4 Fig) we generated for the Y’s from each of these four reads we can conclude that the source of the repeating Y’s are from four different telomeres.

**Figure 5:**
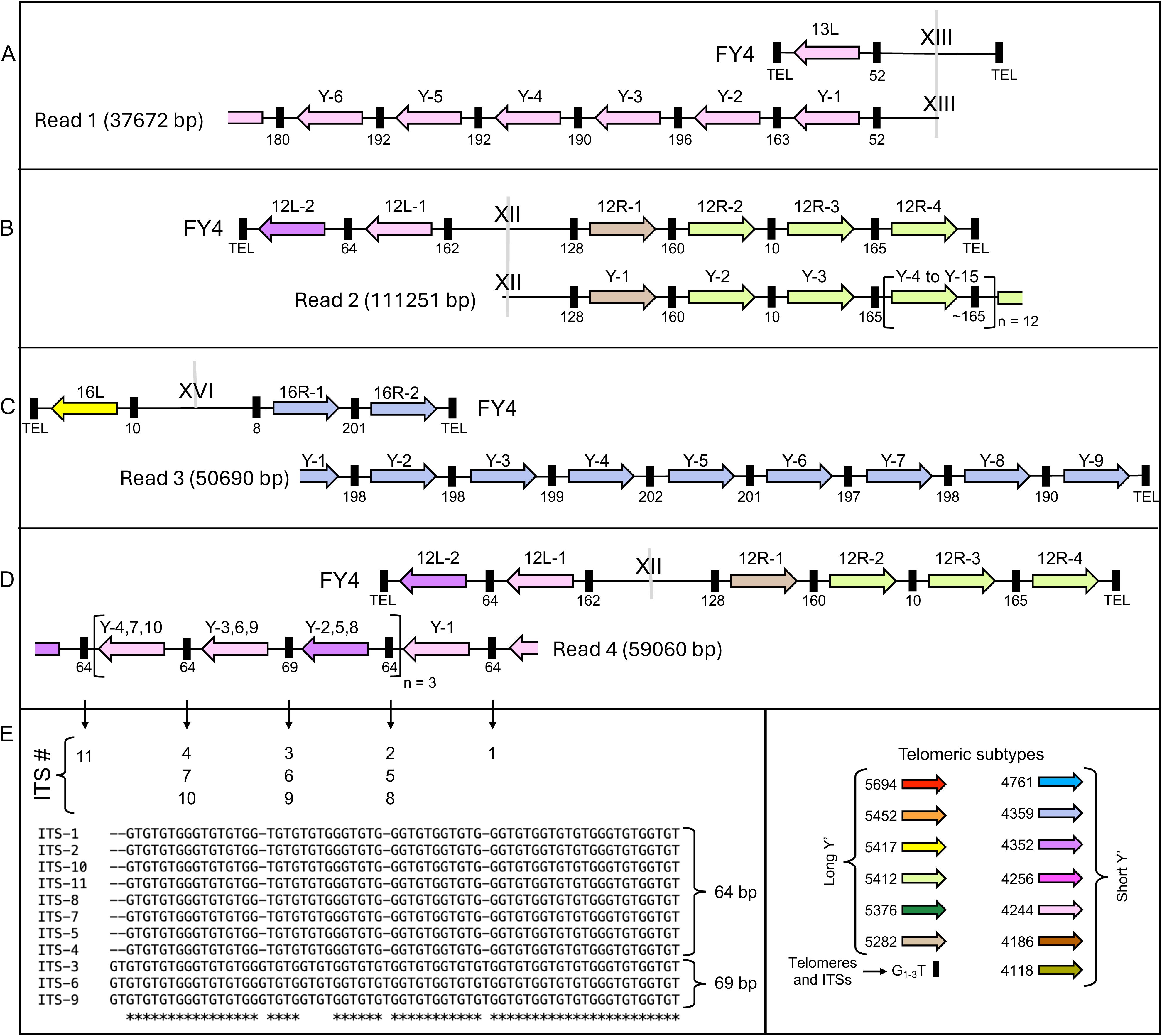
Tandem arrays of Y’s and ITSs are homogeneous and are derived from the resident telomere. In addition to junctions involved in *SUL1* amplification, another major form of junction we identified in *yku70ι1*-A involved tandem Y’ elements separated by ITSs of homogeneous lengths. A-D) Individual Nanopore reads 1 (37672 bp), 2 (111251 bp), 3 (50690 bp) and 4 (59060 bp) are illustrated along with the Y’s from the parental chromosomes for comparison. Square brackets in B and D: sequences that were repeated *n* times as tandem blocks. Multiple alignments of the ITSs in Reads 1-3 (S5A-D Fig) and Read 4 (E) indicate they are unique to each amplicon and homogeneous across the Y’ array. The identities of Y’s (bottom right) were derived by multiple alignments and phylogenetic trees (see S4A-D Fig).

Not only are the contiguous Y’s homogeneous, the ITSs also appear to be related.

The similarity is reflected in the lengths of the ITS (Fig 5A-D) but is also confirmed by sequence alignments (Fig 5E, S5 Fig). There is striking sequence similarity within the groups of tandem ITSs (S5A,B,C Fig) but each form clear clades in the phylogenetic analysis (S5D Fig). From these analyses, we conclude that there is no obvious preference for which Y’ is amplified; however, the resident Y’ and its adjacent telomere appear to act as the seed for generating homogeneity in the tandem array of Y’ and ITSs.

### Model for formation of telomeric tandem arrays

While unequal sister chromatid exchange is an obvious mechanism for producing tandem sequence arrays, the short telomeric tracts at the sites of both *SUL1* and Y’ amplicons in the *ars228* and *yku70Δ*-A and-B mutants suggest a mechanism that involves microhomology such as microhomology-mediated break induced replication (mmBIR). As the name suggests, a break is the initiating event. Telomeres are not usually seen as DNA breaks as they are protected by protein complexes that shield them from exonucleases and end-recognizing machinery that would initiate DNA repair.

However, in the *yku70Δ* mutant, telomeres have lost a major protective protein and fail to efficiently recruit telomerase. As a result, the G_1-3_T telomeric tracts undergo drastic shortening (from a mean of 539 bp in the wild-type to 204 bp in the *yku70Δ* parent; our Oxford Nanopore data). It is the short, unprotected 3’ overhangs that we propose are being recognized as “breaks” and are initiating an mmBIR process.

In Fig 6A we outline an mmBIR event initiated by an unprotected telomere. It is the proposed mechanism for generating type-I survivors in telomerase null mutants (reviewed in [11, 12]) would be so similar, or that the intervening ITSs would be related. With these and other concerns in mind, we propose a version of mmBIR that occurs “*in cis*” (Fig 6B). In summarizing the possible structural changes that could arise from mmBIR, Hastings et al. [24] describe rolling circle replication as a consequence of template switching *in cis* after replication fork collapse. Here, we are suggesting that the unprotected right telomere of chromosome II invades an ITS lying centromere proximal of *SUL1* to initiate conservative replication. When the migrating bubble reaches the initial site of telomere invasion, it now encounters the extended ITS created by that first invasion and continues through the junction sequence into the newly appended repeat. There is technically no end in sight as the BIR bubble continues to re-replicate the sequences just added. One way in which replication could terminate is for the migrating bubble to encounter a single stranded nick on the C_1-3_A strand. This model explains why the *SUL1* amplicons are preferentially found on chrII, why the ITS sequences in tandem repeats are identical (or nearly so), why amplicons suddenly appear with five or more repeats (as in *yku70Δ*-A, Fig 2A,B) and why tandem Y’s are often identical and related to the resident Y’ instead of reflecting the natural Y’ variation seen across the genome.

**Figure 6:**
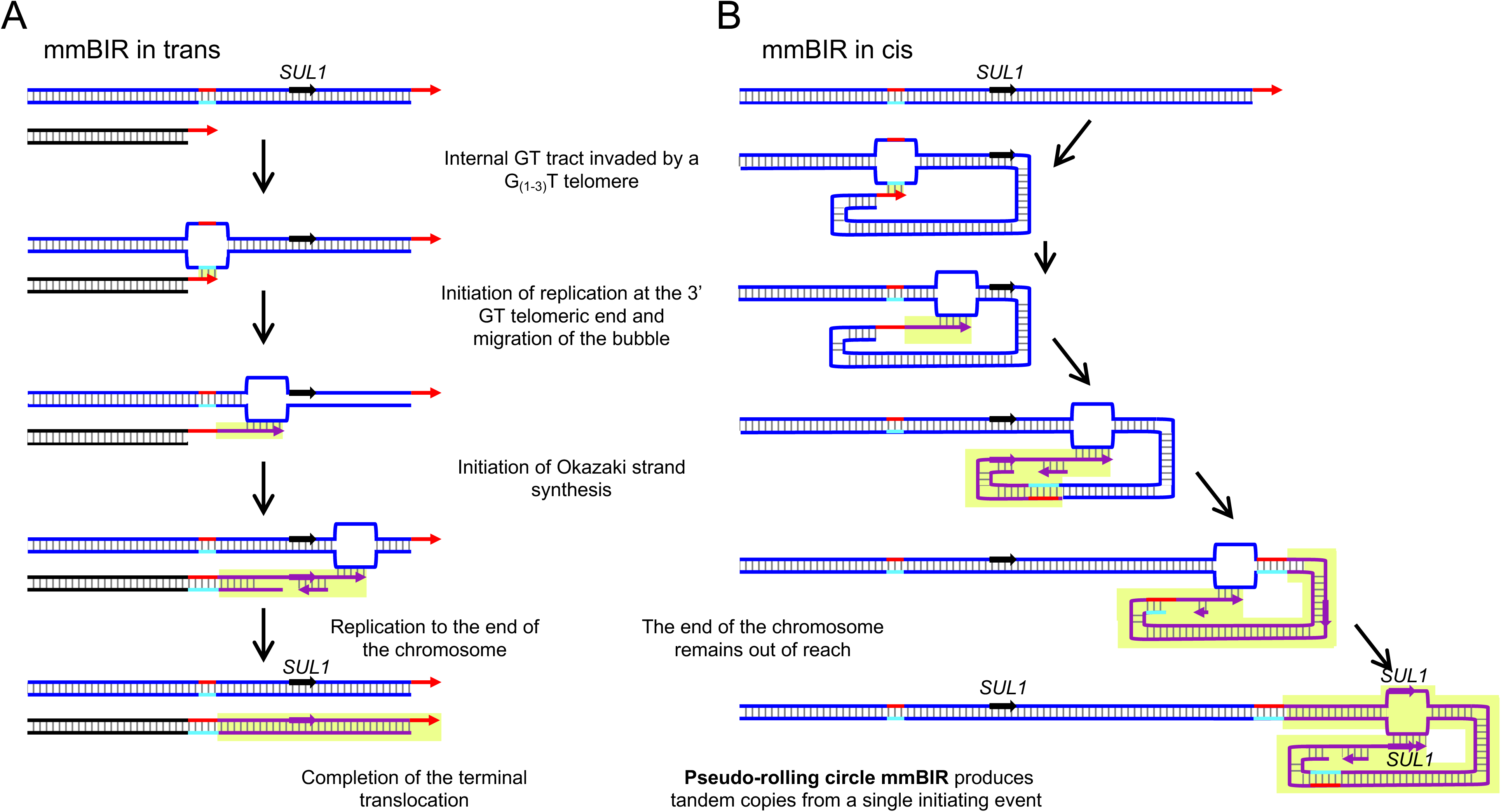
Models that account for tandem telomeric addition. The short stretches of G_1-3_T sequences (ITSs) at the sites of both *SUL1* and Y’ amplicons in the *yku70ι1* mutant suggest a form of break induced replication that is initiated by microhomology (microhomology mediated break induced replication—mmBIR). A) mmBIR *in trans*: When an unprotected telomere (red arrow) invades a sister chromatid, homologue or a non-homologous chromosome at an ITS (red-cyan duplex) the BIR machinery initiates conservative replication that proceeds to the end of the chromosome through a migrating replication bubble. This mechanism can explain the production of a single tandem copy if the invading telomere is from the sister chromatid, or can explain translocations of a Y’ or a *SUL1* fragment when a chromosome other than Chr II-R provides the invading telomere. B) mmBIR *in cis*: When the unprotected telomere invades an ITS on the same chromosome arm it also creates the second copy of *SUL1* and adjacent sequences. However, when the bubble reaches the site of the old telomere, it now encounters the junction created by the invasion and simply copies it and the appended sequences a second, third, or more times. While there is technically no circular DNA involved in this mechanism, it is similar to rolling-circle replication identified in some bacterial plasmids and the yeast 2-micron plasmid during amplification [33]. Hence, we refer to this form of mmBIR as “pseudo rolling-circle mmBIR”.

### Deletion of genes involved in DNA metabolism affects the mode of *SUL1* amplification

The results from *ars228ι1* and *yku80ι1* chemostats suggested that the various pathways of *SUL1* amplification are under genetic control and that mutations in specific genes reduce the incidence of ODIRA events. We expanded our sulfate-limited chemostat analysis to include mutant strains with deletions in genes involved in DNA metabolism (results for nine of the mutants are shown in Fig 7 and S6 Fig). We ran two (or four) independent chemostats for each of the mutants, and in each case, we examined karyotypic changes occurring during chemostat growth by CHEF gels combined with hybridization to a *SUL1* probe (S6 Fig) and performed aCGH on the population from the final day of growth (day 33-35; Fig 7). One of the mutants (*exo1Δ*) failed to amplify *SUL1* or any other unique genomic region. Most populations appeared to have a single amplicon present on the last day. To confirm the form of the amplicons, we had the populations sequenced using Oxford Nanopore long reads and discovered that, in addition to inverted amplicons, there were direct amplicons, ones where the amplified region extended through the right telomere, two translocation events and one extrachromosomal, inverted linear fragment where the junction was composed of sequences from the endogenous 2-micron plasmid (Fig 7, cyan, purple, red, orange bars, green respectively; S1 Table). Only the two chemostats of the *rad5Δ* mutant (Fig 7) produced exclusively inverted repeats. The telomeric amplicons in the chemostats of *msh2Δ, sae2Δ, exo1Δ sgf73Δ* and *pol32Δ* had similar junctions with expanded ITSs (S1 Table).

**Figure 7:**
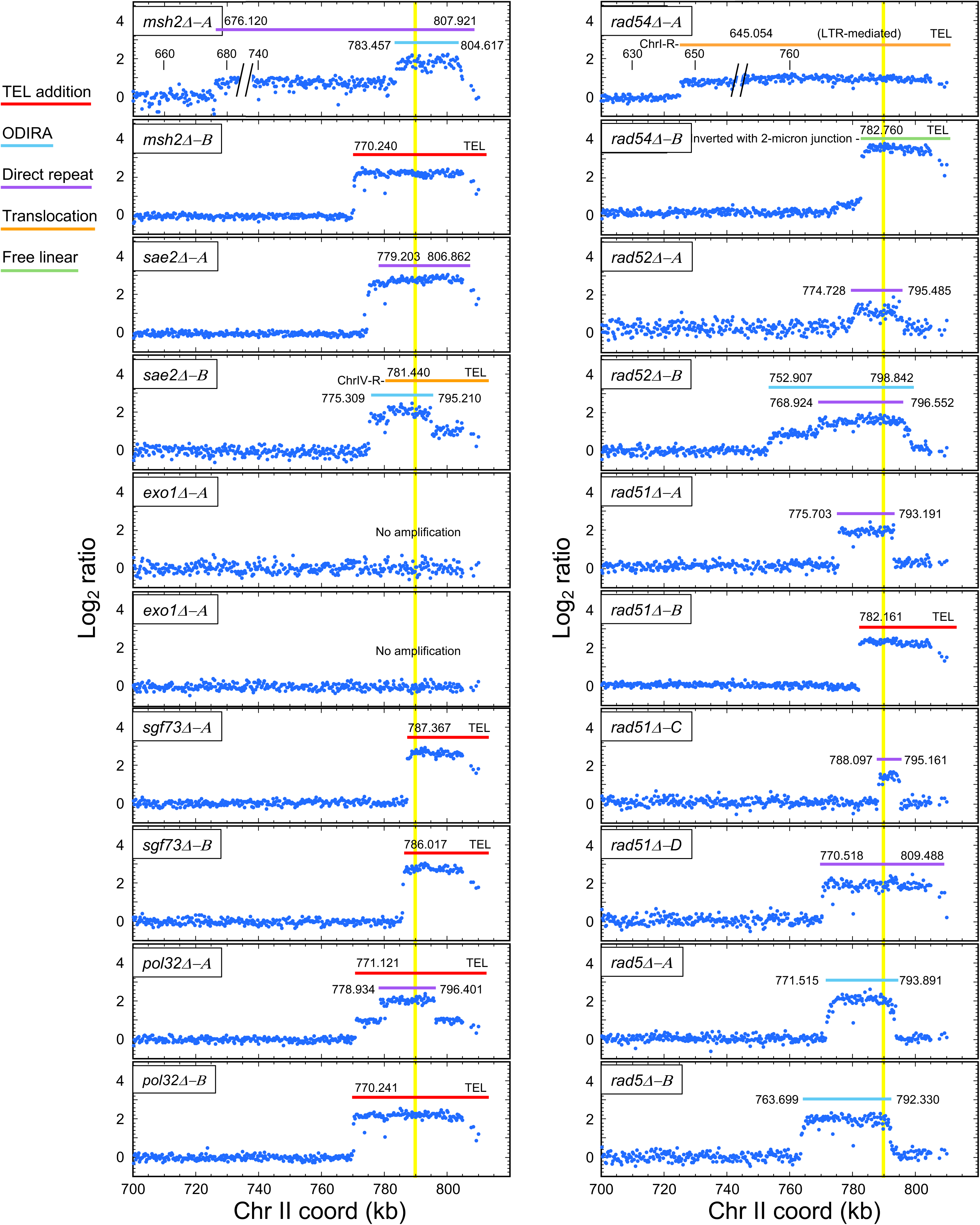
Characterization of chemostat populations of nine gene knockouts by aCGH. Population samples from the final day of chemostats with nine different gene knockouts were examined by array CGH. With the exception of *exo1Δ*, each population generated one or more *SUL1* amplicons (*SUL1* is marked by the vertical yellow bar). The amplified regions were mostly restricted to the terminal 50 kb (exceptions S801 and S1009) and most cultures had resolved to a single predominant amplicon by the end of the experiment (except for chemostats S807, S802, S607 and S606). All junctions were precisely identified by Oxford Nanopore sequencing (see S1 Table) and are indicated by chrII coordinates (in bp). Colored bars indicate amplification through the telomere (red), inverted (ODIRA) amplicons (cyan), direct repeats (purple), translocations (orange) and free linear fragments (green).

To quantify the effects of different mutations on the ODIRA pathway of gene amplification, we moved four of the mutations (*yku70Δ*, *sae2Δ, rad5Δ*, and *rad54Δ)* into our test strain (s2-1) that harbors a split-*URA3* cassette (S7A Fig; [2]). The position of the two overlapping partial *ura3* fragments generates a specific chromosomal rearrangement when a *URA3* gene is produced by an ODIRA-like event. We chose these mutants because they spanned the range of only ODIRA events (*rad5Δ*), only terminal events (*yku70Δ*), and a mixture of four forms of amplification (*sae2Δ* and *rad54Δ*). In wild type cells, the frequency of ODIRA events is 22% (11/50 ura+ clones; [2]). ODIRA events were reduced in three of the mutants (*rad5Δ*, *yku70Δ, rad54Δ*) and increased slightly in the *sae2Δ* mutant (S7B Fig).

We cannot say definitively, without further work, how the different mutant genes we tested might be participating in the ODIRA or mmBIR pathways. For example, we cannot say whether Yku70 or Sgf73 (whose deletions produced only telomeric *SUL1* amplicons, Fig 2 and Fig 7) are required for the ODIRA mechanism or their absence makes telomeric invasion more likely, or perhaps both. That would require measuring absolute rates of these two mechanisms in the chemostats—something that we are currently unable to do experimentally.

### Test of the *cis*-mmBIR mechanism for tandem addition of subtelomeric sequences

The genetic requirements for mmBIR are well established (reviewed in [11, 12]).

In particular, mmBIR is dependent on the DNA polymerase that contains a subunit encoded by *POL32*. Since the two *pol32Δ* chemostats (Fig 7) both produced amplicons of *SUL1* that were terminally duplicated it appears that our model cannot be correct. However, terminal duplications could also be created by non-allelic homologous recombination. To determine to what extent Pol32 was involved in terminal duplications we tested a *pol32Δ yku70Δ* double mutant. To simplify the analysis and avoid the multiple sulfate-limited chemostats that would be required, we employed a batch culture protocol to ask whether simply passaging strains would produce the same unstable Y’ phenotype. We performed three technical replicates for each of four strains (wild type, *yku70Δ*, *pol32Δ*, and *pol32Δ yku70Δ*), growing them for 22 consecutive days (∼130 generations) in the same sulfate-limiting medium that we had used in the chemostats. Because batch culture cannot reproduce the same selective conditions for *SUL1* amplification that a chemostat can, we did not expect to see *SUL1* amplification (and we recovered none) but we were able to reproduce the Y’ instability.

At the end of the 22 days of culture, we isolated 12 clones from each of the 12 cultures and performed CHEF gels and Y’ hybridizations of the Southern blots (Fig 8A-D; S8 Fig). It is clear by inspecting the Y’ autoradiograms of the *yku70Δ* Southern blots that there are no parental karyotypes remaining and that many of the chromosomes with Y’ additions have a very significant increase in the amount of Y’ hybridization on most of the altered chromosomes relative to the hybridization levels on chromosomes in the other three strains (Fig 8A-D; S8 Fig). We identified a total of 7 of 34 clones with Y’ additions in the wild type strain and 34 of 34 in the *yku70Δ* strain (S8 Fig). This result clearly indicated that continuous passaging of the *yku70Δ* strain was sufficient to result in Y’ amplification. In contrast, among the 35 unique clones of the *pol32Δ yku70Δ* strain only nine had an altered chromosome size, a frequency that was identical to the number of clones we found in the *pol32Δ* strain (S8 Fig). Thus, deleting *POL32* strongly reduces the Y’ amplifications caused by deletion of *YKU70*, underscoring the important role that Pol32 plays, specifically, in the pathway that occurs when telomeres are destabilized by the absence of the Ku heterodimer. Nevertheless, the fact that there were non-zero numbers of terminal Y’ amplification events in the absence of Pol32 confirms that there is not a single pathway to generate terminal duplications.

**Figure 8:**
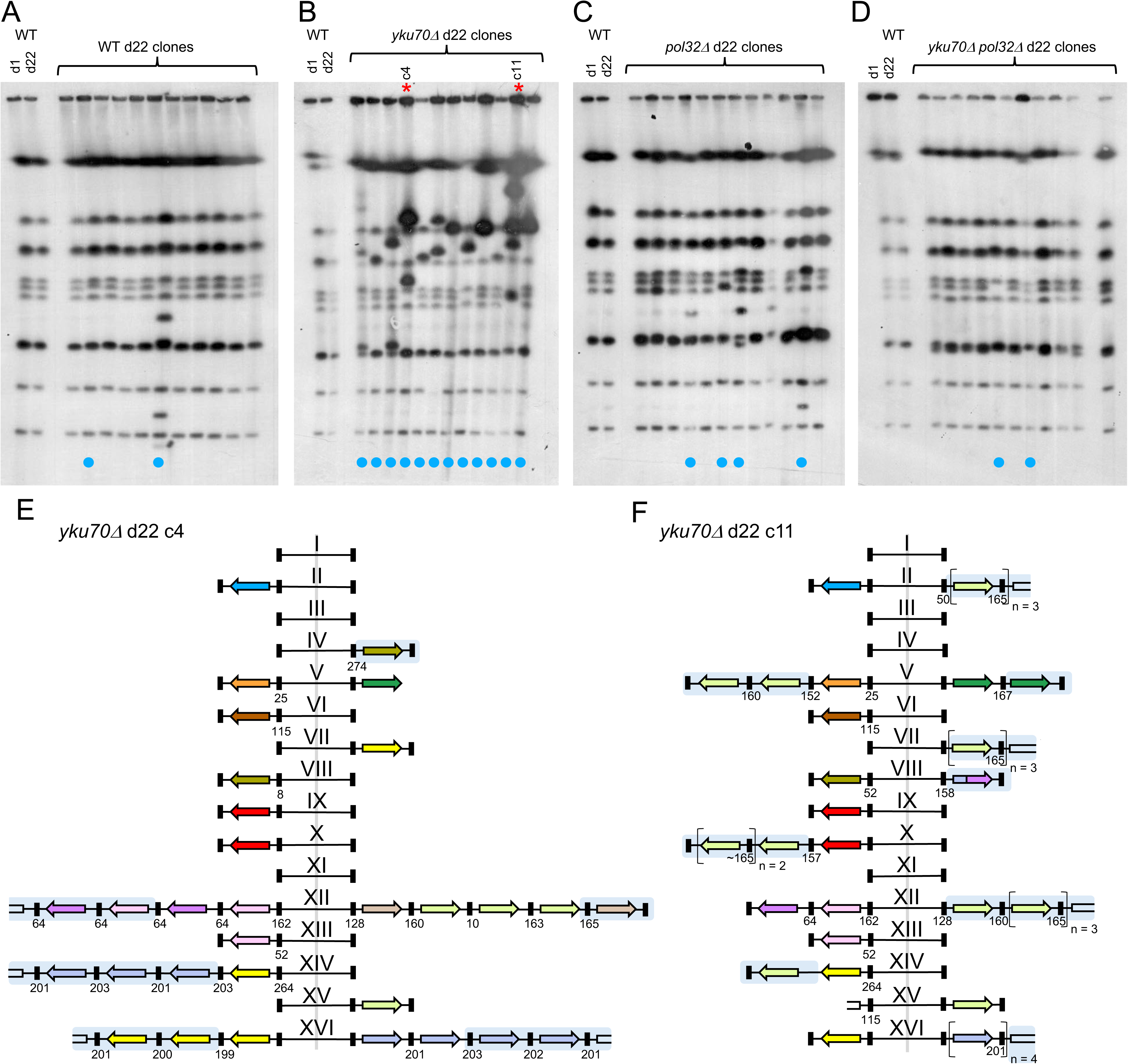
**Batch culture evolution of *yku70****Δ* **confirms that Y’ amplification events are decreased when combined with a *pol32****Δ* **mutation** Four strains (wild type, *yku70Δ*, *pol32Δ,* and *yku70Δ pol32Δ*) were serially transferred for 22 days (∼140 generations). On day 22, 11-12 colonies from each culture were examined by CHEF gel and Southern hybridization with a Y’ probe (A-D). Clones with altered Y’ hybridization are indicated by cyan circles. Two clones from *yku70Δ* (B; c4 and c11, indicated by the red asterisks) were sequenced with Oxford Nanopore Technology and their telomeric regions examined for amplification. The chromosomal assignments of Y’ telomeres in clones 4 and 11 are diagrammed in E and F. Structural alterations are highlighted by blue shading. The phylogenetic analysis of the Y’s from these two clones are shown in S9A,B Fig. Southern blots of CHEF gels from twelve clones from each of two additional technical replicates are shown in S8 Fig. The telomere maps and phylogenetic analysis of the three additional clones are given in S9C,D,E Fig.

To confirm the structure of the Y’ amplicons that arose in the batch culture of the *yku70Δ* strain we had five clones sequenced using Oxford Nanopore technology (c4, c11, c6, c12 and c10; Fig 8E,F; S8 Fig; S9 Fig). Each had tandem additions of homogeneous Y’s that were related to the resident Y’ and were separated by ITSs of similar size. Additional features we identified in the clones were the secondary transfer of amplified Y’ arrays to other chromosomes, presumably through homologous recombination (e.g., c4 chr XIV-L; c11 chr X-L; Fig 8E,F) and two hybrid Y’s with the 5’ end of one chromosome and the 3’ end of another chromosome (Fig 8F; S9B Fig; S9D Fig). We also found examples where a Y’ had been added to the end of a chromosome that had previously lacked a Y’ (Fig 8F chrII-R; S9D Fig chrII-R and chrXI-L), indicating that mmBIR occurs *in trans* at a reduced frequency.

### Terminal amplification in the human genome explained by mmBIR *in cis*

The Telomere-to-Telomere (T-2-T) human genome assembly has added important sequence details in repeated parts of the genome, including refinements at telomeres. In a scan of the human T-2-T genome (https://genome.ucsc.edu) we focused on the recent addition to the reference genome of ∼160 kb of sequences at the chromosome 18p telomere. In that region are four ∼45 kb repeats that include several annotated genes (including genes for tubulin variants and for LINC RNAs) (Fig. 9A; S10 Fig). Between the direct repeating units are internal stretches of the CCCATT telomeric repeats that are strikingly similar to each other in size (∼650 bp) but significantly shorter than the average terminal telomeric CCCATT tract (Fig 9B; S10 Fig). The distribution of sizes is comparable to the distribution of telomeric lengths in yeast and the ITSs in the Y’ tandem arrays we found in the *yku70Δ* mutant (Fig 9C). This pattern at chr18p is similar in size and structure to terminal *SUL1* amplicons we found arising in the yeast *yku70Δ* strain and suggests the possibility that this human amplification event may have occurred by mmBIR pseudo-rolling-circle replication during a period of telomere stress.

**Figure 9:**
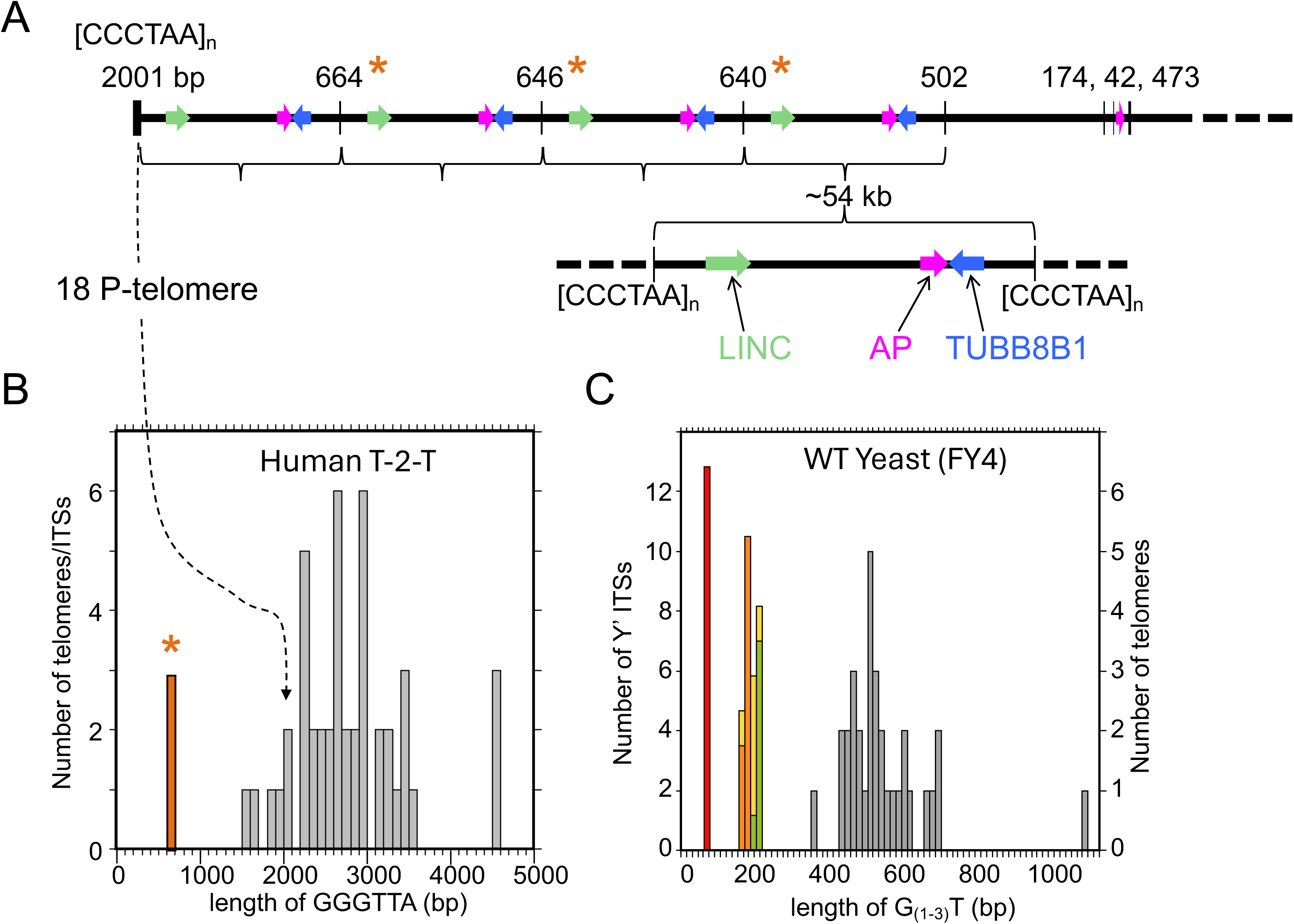
Tandem telomeric amplification on human chromosome 18p. Human telomeres are capped by the sequence TTAGGG/CCCTAA, and as in yeast, these repeats are found throughout the genome as ITSs. A) A simplified drawing of ∼300 kb of telomere adjacent sequence of Chr18p from the T-to-T human genome sequence illustrates the telomere and several adjacent ITSs (red asterisks). Several genetic landmarks (LINC, AP and TUBB8B1) illustrate the repeated nature of this region and separating each repeat is an ITS of relative uniform size (∼650 bp). B) A histogram of telomere tract length was compiled from the T-to-T data for the 23 chromosomes and compared to the ITS lengths from Chr18p, highlighted in red in the histogram. (A screen shot of this region taken from the human genome browser is shown in more detail in S10 Fig). C) A similar histogram of telomere lengths in FY4 yeast (gray) and the ITSs from the four tandem Y’s from Fig 5 (red, orange, yellow and green) suggest a common mechanism for terminal amplification events from yeast to humans.

The initial step in our proposal for mmBIR *in cis* shares structural similarity to the T-loops found in mammalian chromosomes [25]. These T-loops form by invasion of the single stranded G-rich strand into the upstream duplex tract of telomeric repeats and can range from 1 to 25 kb in size. They are formed and stabilized by the Shelterin complex and are thought to protect chromosome ends from degradation and/or recombination [25]. The authors speculated on the potential for the invading 3’ end to initiate replication, and it is now commonly thought that replication from this imbedded 3’end is responsible for the lengthening of the G-rich telomeric tract in ALT type-II survivors. More recent work has shown that invasion of the 3’ end of the G-rich strand can also associate more distally with ITSs [26], but no role for replication priming has been proposed. More long read sequencing, particularly of telomere-adjacent regions, in individuals or tumors suspected of copy number disorders will potentially determine whether there is a role for telomere associated mmBIR *in cis* in human disorders.

## CONCLUSIONS

Selection is a powerful thing “but life finds a way” [27Context for the quote: “Because the history of evolution is that life escapes all barriers. Life breaks free. Life expands to new territories. Painfully, perhaps even dangerously. But life finds a way.”]. In chemostats limiting for an essential nutrient, yeast reveals that there are multiple pathways for amplifying genes that provide a growth advantage. By interfering with the most common mechanism used to amplify the *SUL1* gene (ODIRA) we have shown that cells co-opt a mechanism (mmBIR) involved in cell survival when telomerase is deleted (ALT Type-1) to increase *SUL1* copy number. We have also shown that ALT is not limited to telomerase-null mutants but can occur in wild type cells and other mutants that alter DNA maintenance. In the process of analyzing these new amplicon variants, we have refined the understanding of the ALT mechanism by proposing that the strand invasion predominantly occurs *in cis* through the formation of a structure similar to naturally occurring T-loops found at the ends of mammalian chromosomes. It remains to be seen to what extent this mechanism of gene amplification contributes to *de novo* mutations in human evolution, development disorders, aging or cancer.

## ACKNOWLEDGMENTS

MJD thanks Grant Brown and Barry Dion for early experiments exploring how DNA repair mutants adapt in chemostat culture and Alex White for technical assistance with chemostats. BJB and MKR thank Bob Waterston for the gift of two additional Bio-Rad CHEF gel apparatuses and the many dedicated undergraduate student helpers who helped keep our lab running during the course of this work.

## MATERIALS and METHODS

### Stains and culture conditions

All strains in this study were derived from FY4 or its close relative BY4741. The split-*URA3* strain, s2-1 [2], was used to quantify the effect of *yku70Δ*, *sae2Δ* and *rad5Δ* on the frequency of ODIRA amplification events. Medium and culture conditions for the sulfate-limited chemostats are as described [1, 28]. Batch culture passaging was carried out in 6 well plates with 5 ml of sulfate-limited medium at 30°C. A 100-fold dilution was made every 24 hours for 22 days and performed in triplicate for each strain. Twelve clones from each culture from day22 were isolated for further analysis. All gene deletion mutants were obtained from the KanMX yeast knock out collection [29, 30] and/or created by transformation using PCR fragments and confirmed by PCR and Sanger sequencing (Eurofins). The *ARS228* deletion was created by scrambling the ARS consensus sequence [4].

### Molecular analysis of clones and cell populations

Periodically, during growth in chemostats or serial transfer, genomic DNA from cell populations was analyzed by CHEF gel electrophoresis [31, reviewed in 32] and Southern blotting [14] to detect karyotypic changes. The population or single clones from the final day of chemostat growth were analyzed by aCGH [2, 14, 31] to identify the segments of the genome that had been amplified. The final population of one chemostat (*yku701′*-A) was sequenced in house using Oxford Nanopore Technology.

Sequences and contigs (S1 Data) of clones or populations from other chemostats, batch culture evolutions and the parental FY4 and *yku701′* parent strains were performed by Plasmidsaurus using Oxford Nanopore Technology (S2 Table). Raw reads from these data were analyzed using BBEdit (Bare Bones Software) and alignment to the reference genome to identify amplification junctions. Short read sequence data for populations of 31 independent sulfate-limited chemostats are from Martin et al., 2024 [2].

### Characterization of *SUL1* and Y’ amplicons

Split reads from Oxford Nanopore sequencing of the population from the *yku701′*-A chemostat were examined manually in IGV to identify *SUL1* amplicon junctions.

Sequence coverage of individual junctions ranged from 1 to 62. (The single split read for the junction at 786.502 kb was present twice on the same fragment as part of a *SUL1* triplication.) Among the longer reads (>∼40 kb) we found evidence for tandem amplicons of 3-4 repeats for five of the junctions. We were also able to identify translocations of the *SUL1* fragment to two other chromosomes.

The split reads from Oxford Nanopore sequencing of *yku70Δ*-A also confirmed the tandem structure of Y’s with intervening ITSs. To determine the identities of the Y’s involved, we used tandem Y’s in a BLAST search and found chr XII (left or right) was always the best match, presumably because chrXII in SGD has tandem Y’s at each end. However, when the same sequence was split into individual Y’s, other telomeres provided the best BLAST match. Because some chromosomal ends are incomplete in SGD we focused on a section of the Y’ region (between 5’ATGGAAATTGAAA and 5’ATGCACTTGCGAGATC) common to all Y’s in SGD and performed multiple alignments (CLUSTALW) and determined phylogenetic relationships (FastTree full) of the Y’s in the reference genome, and compared them to Y’s in FY4 and the *yku70Δ* strains (S3 Fig).

The phylogenetic relationships allowed us to define thirteen Y’ classes (S3A Fig Legend) and their distribution across the sixteen yeast chromosomes (S3A Fig). (Trees generated by PhyML or RAxML had identical topologies.) The Y’ identities on individual chromosome ends for the three strains mostly agree. We used this same strategy of creating multiple alignments and phylogenetic trees to identify the origin of Y’s in the tandem repeats found in our experimental samples. However, for Y’s we compared tandem repeats from single long reads to the *yku70Δ* parent consensus sequences.

Even with the relative high frequency of sequencing errors it was still possible to identify the closest Y’ relative for the individual Y’s in the tandem repeats.

## SUPPLEMENTAL FIGURE LEGENDS

**S1 Figure:**
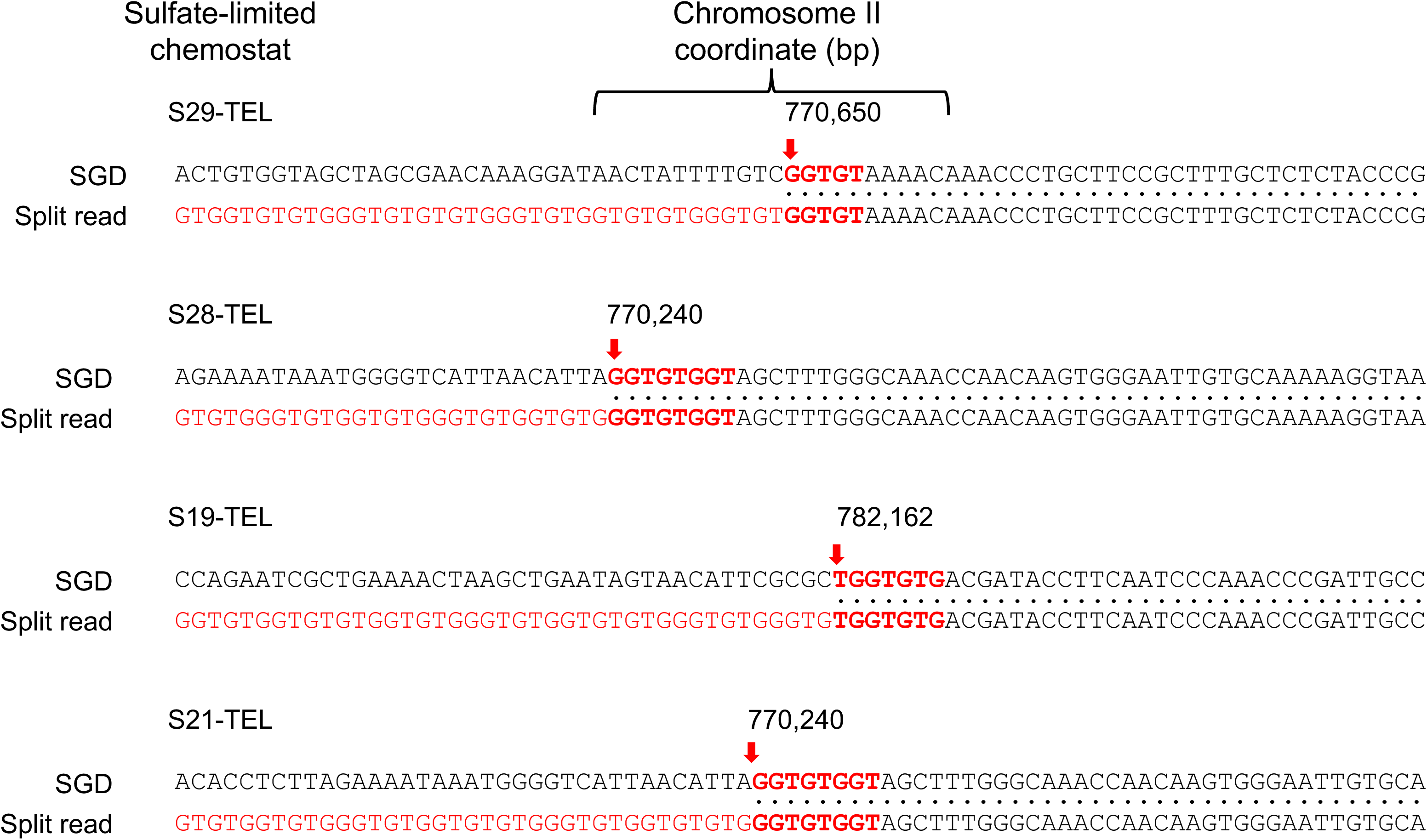
Rare terminal *SUL1* amplicons in wild type yeast. Among 31 sulfate-limited chemostat populations that we analyzed by short read sequencing [2] we discovered four clear examples of a split read between a telomeric sequence and unique sequences on chr II that were centromere proximal to *SUL1*. The junction at 770.240 kb was found in two independent cultures. Each of the reads resulted in an expansion of the resident G_1-3_T sequence tract and is similar to the clone recovered in the *ars228Δ* chemostat. For comparison, in these same 31 evolutions, there were 50 inverted junctions (centromere proximal to *SUL1*) consistent with an ODIRA mechanism of *SUL1* amplification [2].

**S2 Figure:**
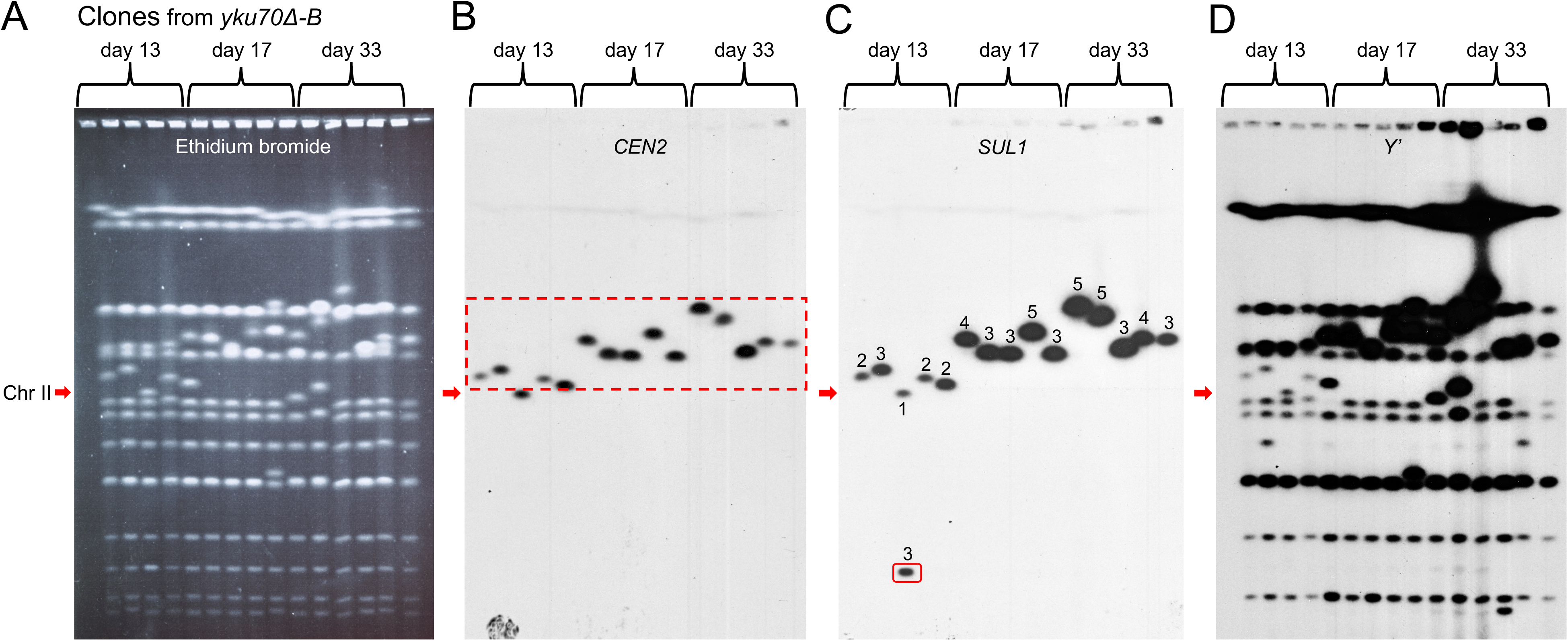
**Y’ amplification in *yku70****Δ***-B is a consequence of continuous culture in the sulfate-limiting chemostats** Fifteen clones from days 13, 17 and 33 of *yku70Δ*-B were isolated and compared to the wild type strain using CHEF gel electrophoresis and ethidium bromide staining (A) and Southern blotting (B,C,D) as described in the legend to Fig 3. These results were consistent with the observations made on clones from *yku70Δ*-A. Dashed red box, clones with an altered size of chr II; solid red box, a clone where *SUL1* has moved to a different chromosome. Numbers above the bands in (C) indicate estimated number of copies of *SUL1*.

**S3 Figure:**
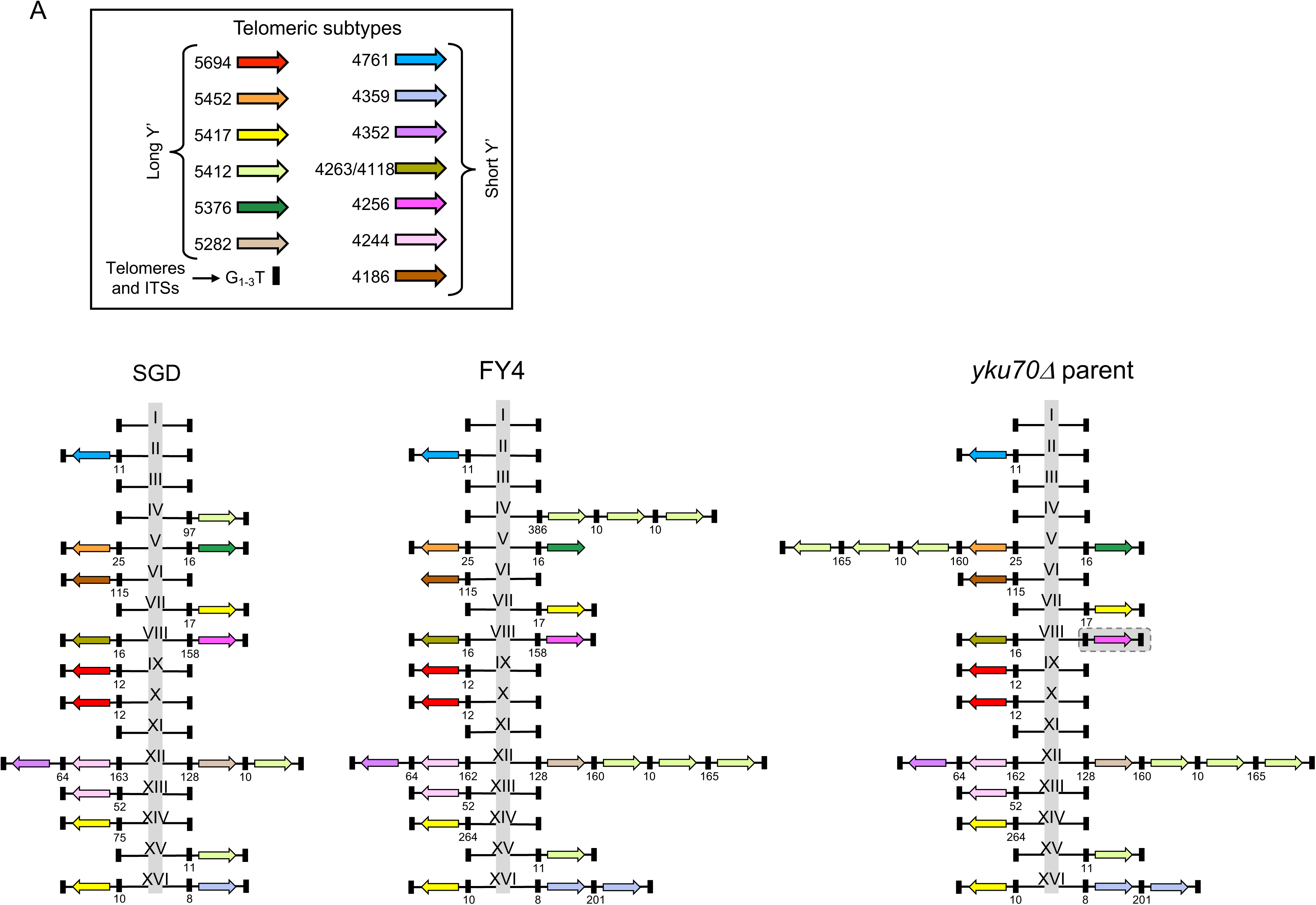
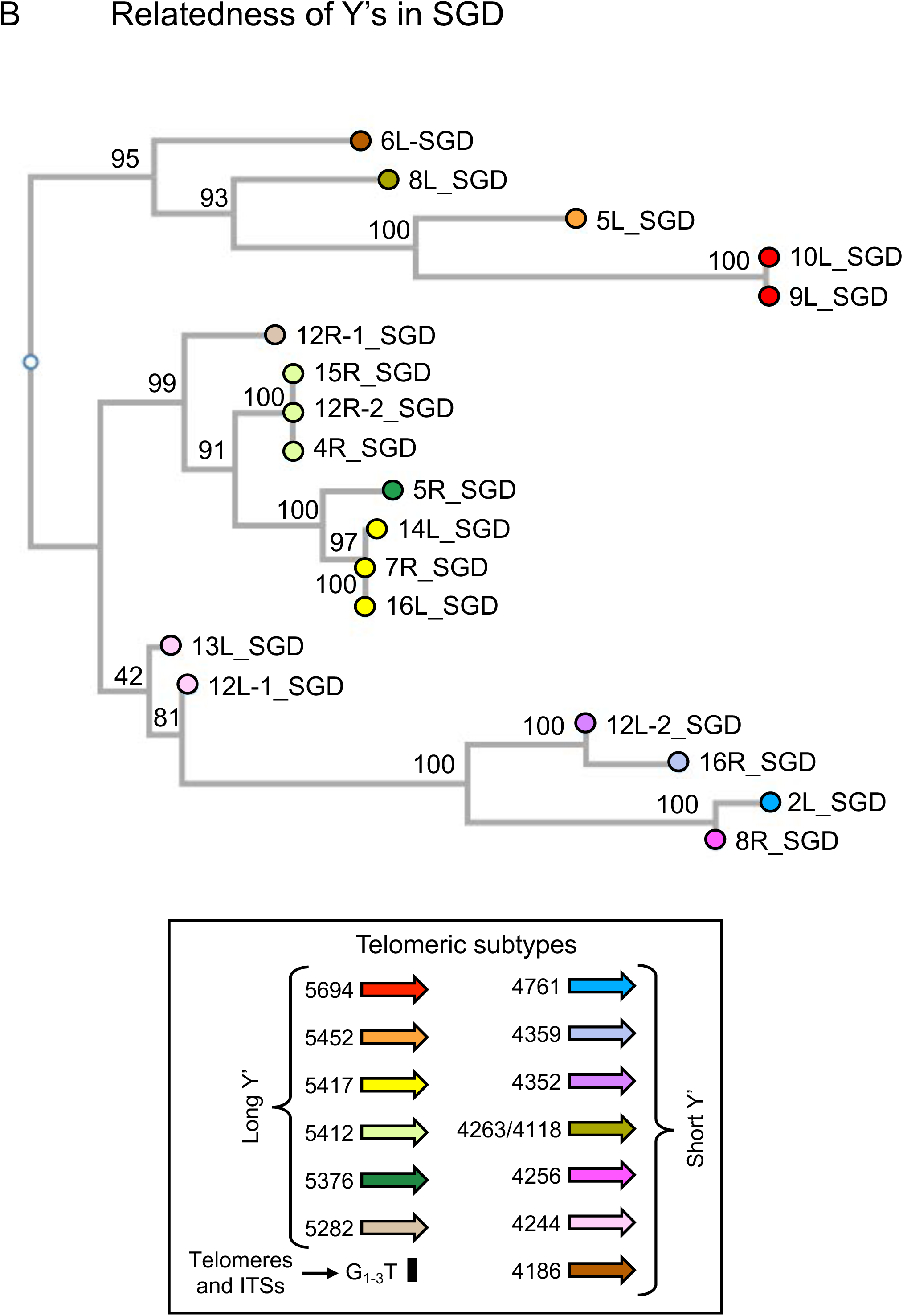
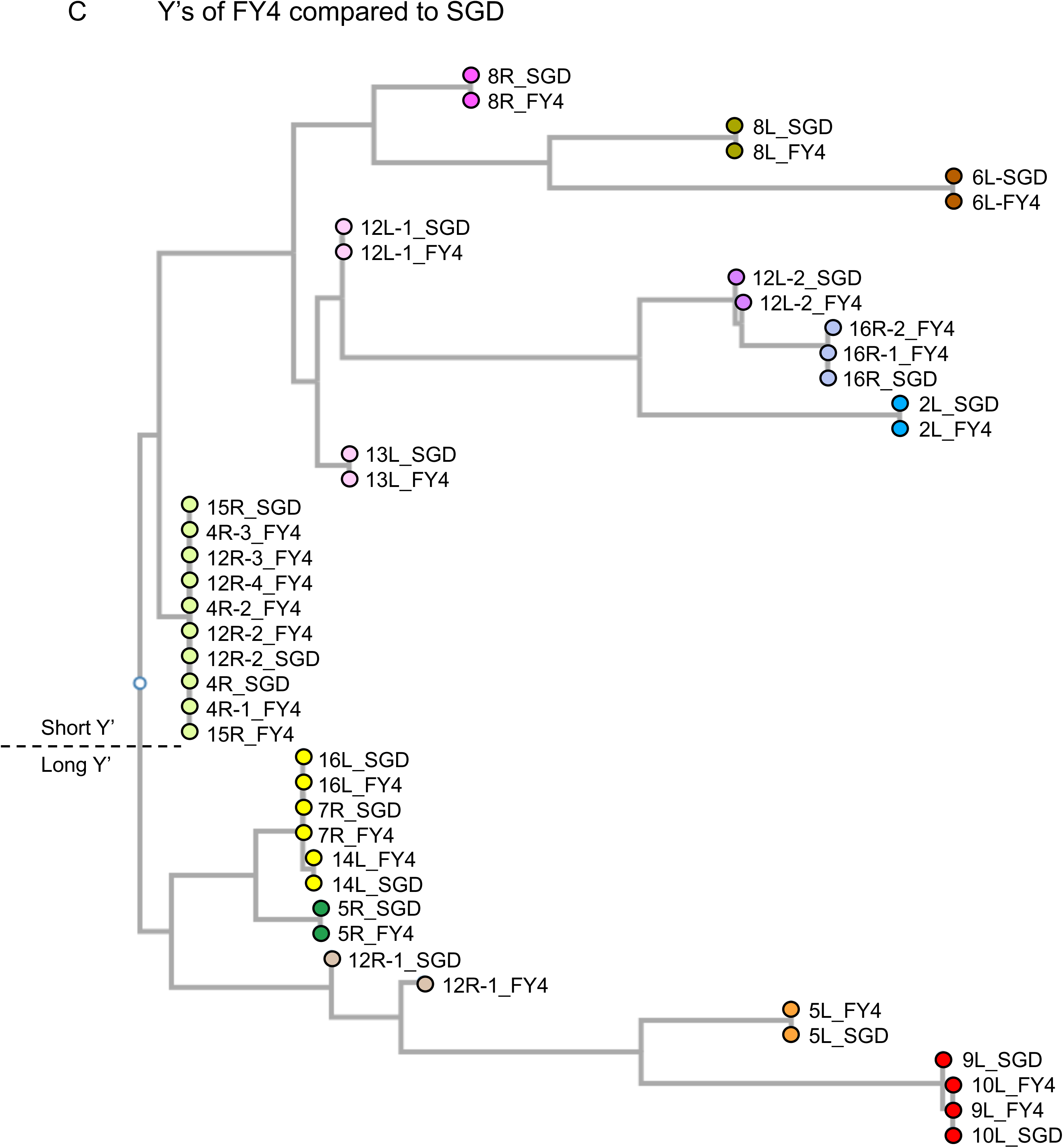
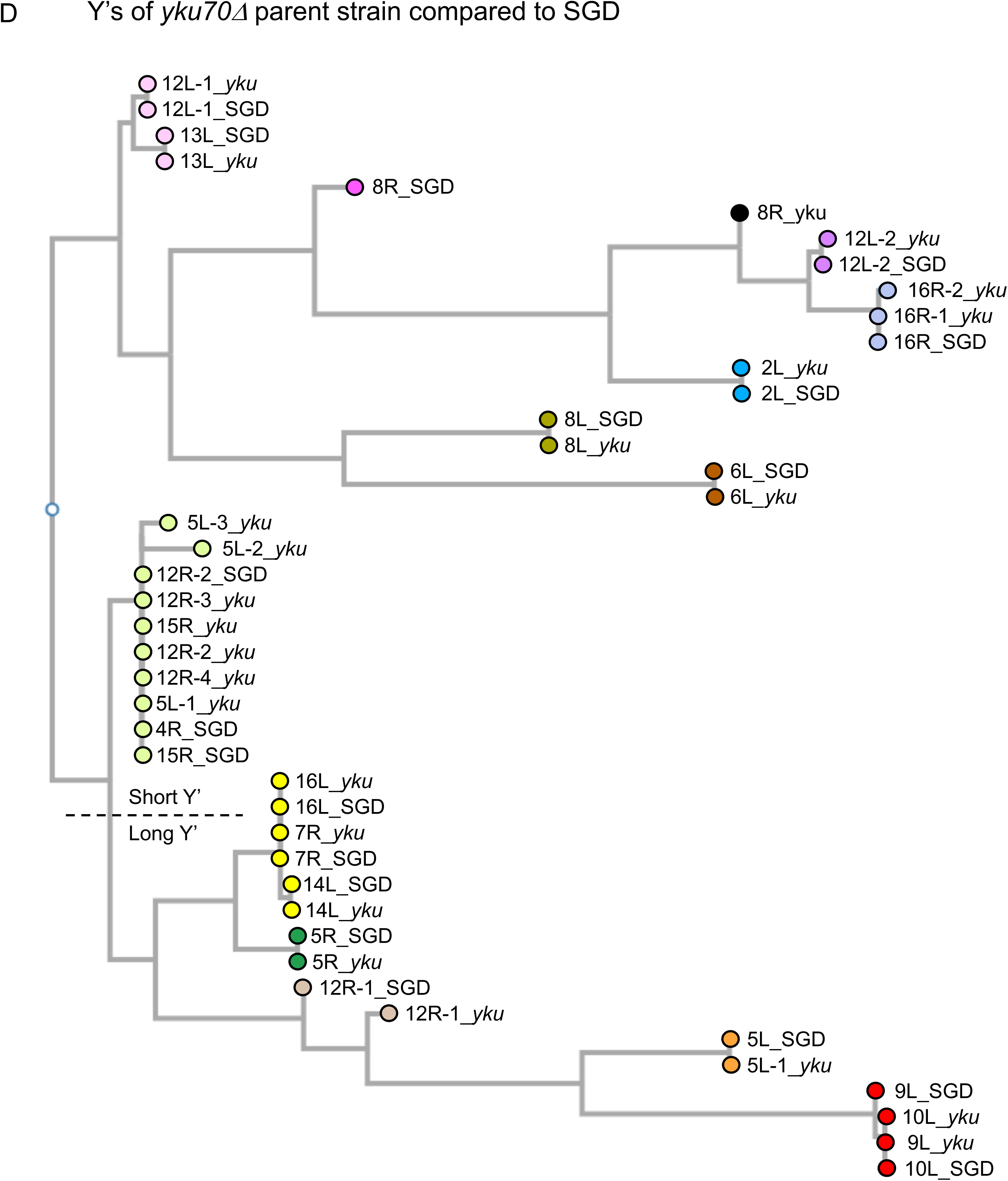
**Phylogenetic relationships of Y’s between the reference genome (SGD) and wild type FY4 and the *yku70****Δ* **parent strain** A) The sixteen chromosomes of the Saccer3 reference genome (SGD) and the two strains used in this work (FY4 and *yku70Δ* parent) are diagrammed to highlight the differences in telomere structures with the unique portions of the chromosomes being reduced to the vertical gray bar that is labeled I through XVI. Note: We are assuming that all chromosomes in SGD end in G_1-3_T/C_1-3_A sequences and are thus included in the illustrations of SGD. Nanopore sequencing confirmed that FY4 and the *yku70Δ* parent chromosomes ended in G_(1-3)_T sequences, with the exception of the right telomere of ChrVIII in the *yku70Δ* parent (shaded rectangle) which ended in repeated sequences upstream of the Y’ in our sequencing. We infer that this Y’ is present because it appears in Nanopore reads from descendants of this strain. B-D) To distinguish individual Y’ subtypes we aligned the sequences of 19 Y’s in the reference genome (defined by boundary sequences of 5’-ATGGAAATTGAAA-3’ and 5’-ATGCACTTGCGAGATC-3’) and created relatedness trees using CLUSTALW and FastFull Tree programs. The bootstrap values and the branch lengths allowed us to identify thirteen distinct Y’ sequences (six long versions and seven short versions; legend). The phylogenetic relationships of Y’ between SGD and FY4 and between SGD and the *yku70Δ* parent indicate nearly perfect agreement.

**S4 Figure:**
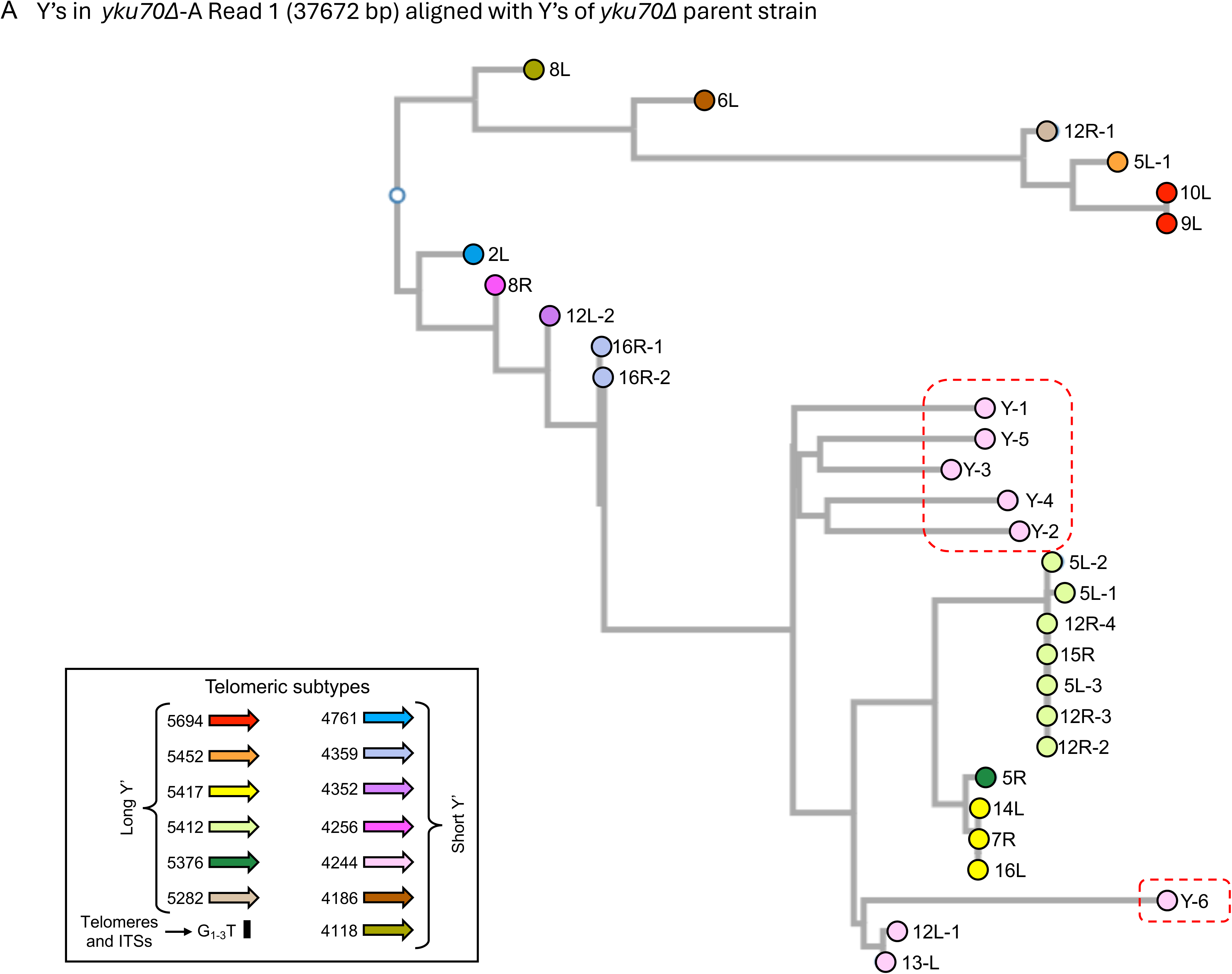
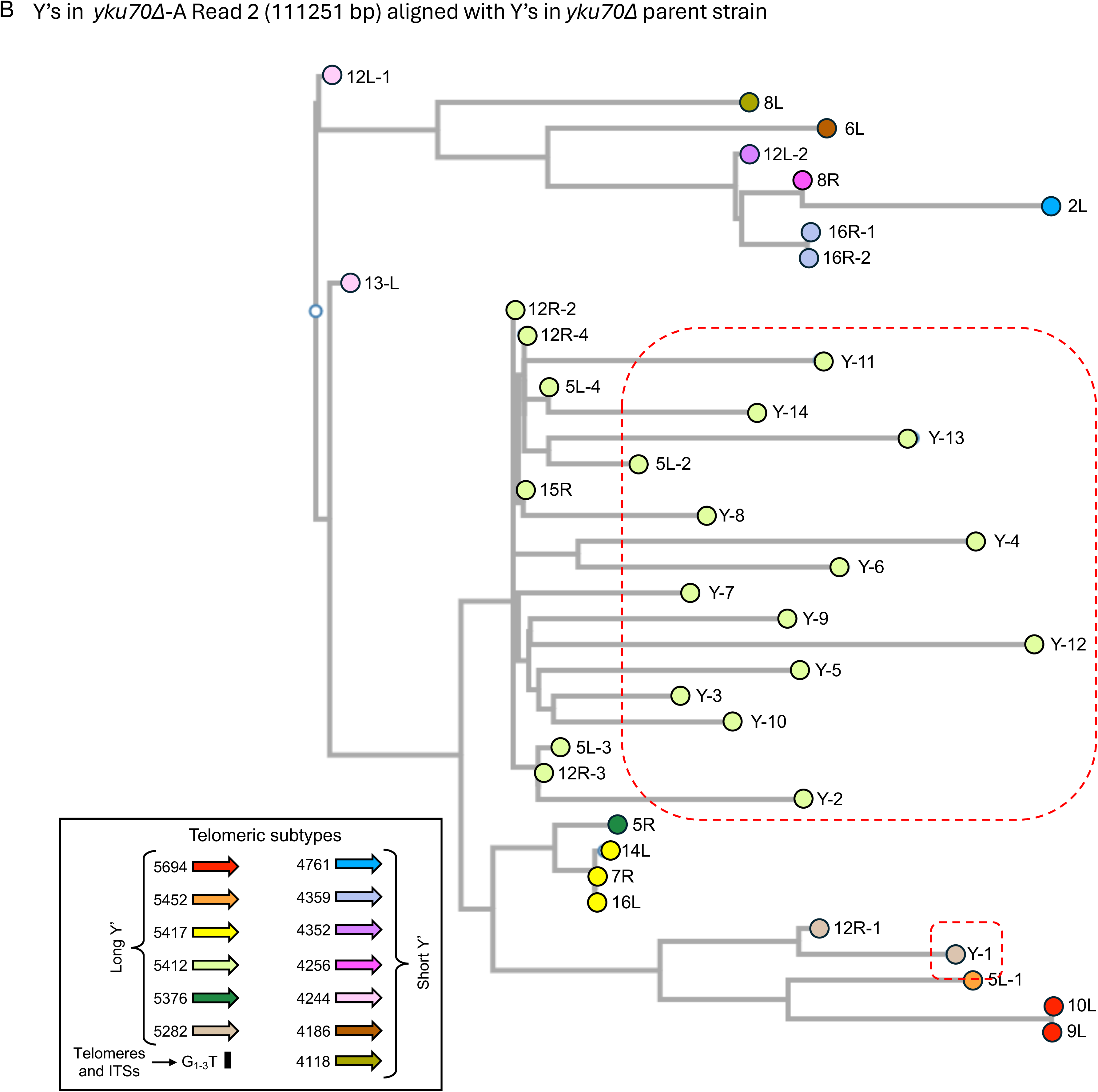
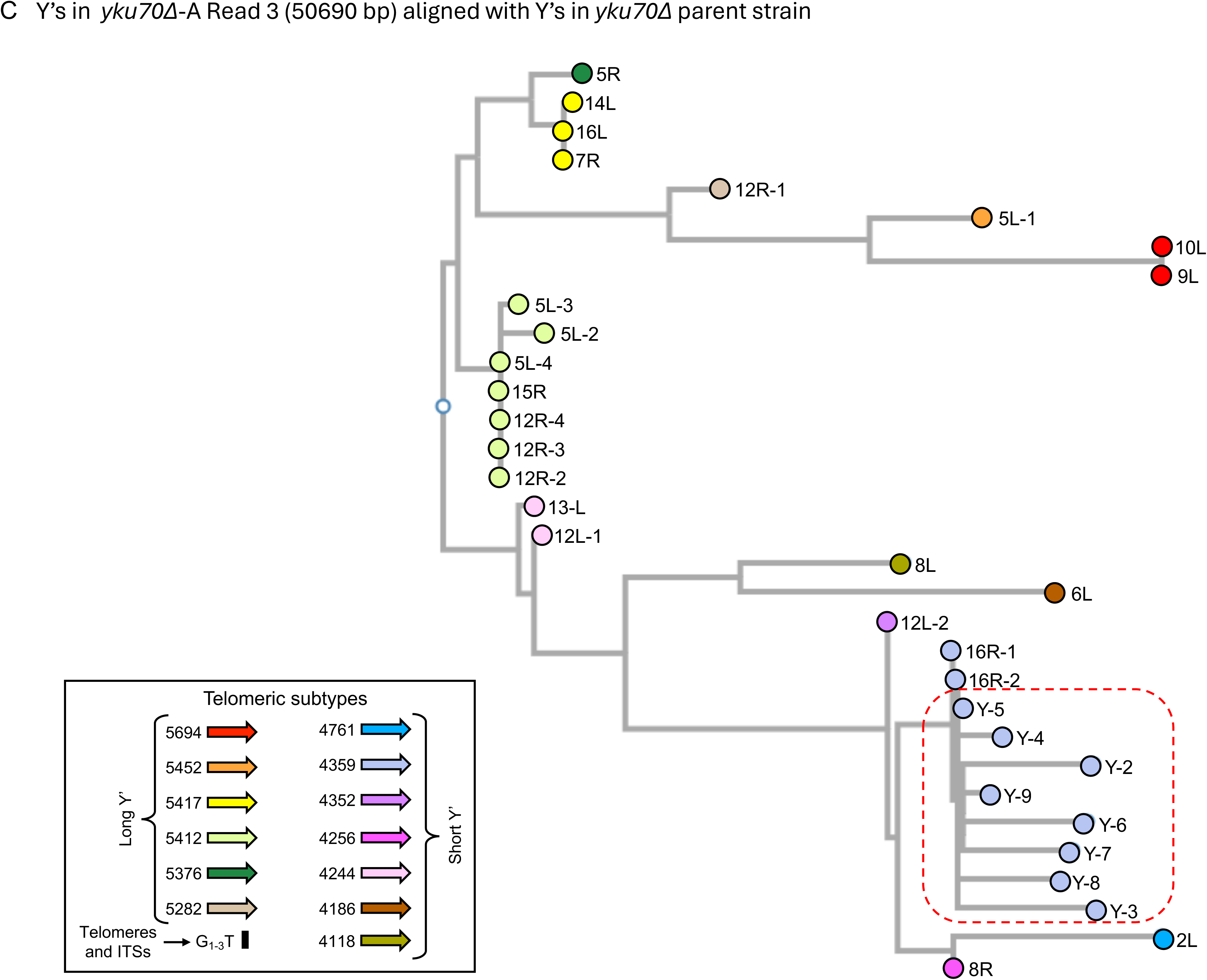
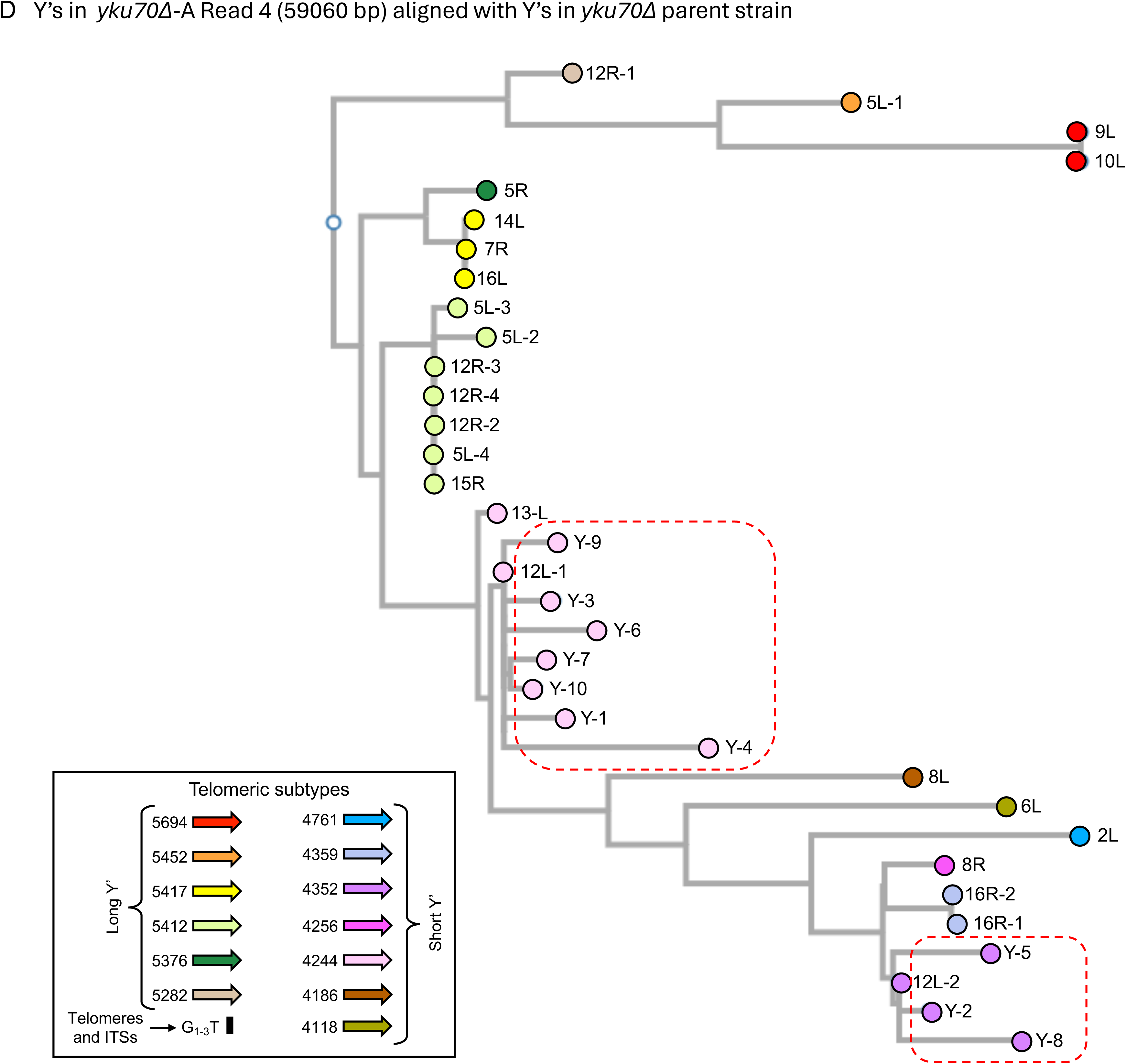
**Phylogenetic trees comparing the *yku70****Δ* **parent with individual Nanopore reads identify the source of Y’s in the amplified tandem arrays** Trimmed Y’s from the Nanopore contigs of the *yku70Δ* parent (day 0) were aligned with the trimmed Y’s from individual Nanopore reads from *yku70Δ*-A: A) Read 1 (37672 bp); B) Read 2 (111251 bp); C) Read 3 (50690 bp); and D) Read 4 (59060 bp). Read 3 had a partial Y-1, which was excluded from the alignment. In each case the amplified Y’s cluster with the parental Y’s (boxed in red dashed line). The divergences from the ancestral sequences are exaggerated by errors produced in sequencing as they represent a single read and not a consensus across multiple reads.

**S5 Figure:**
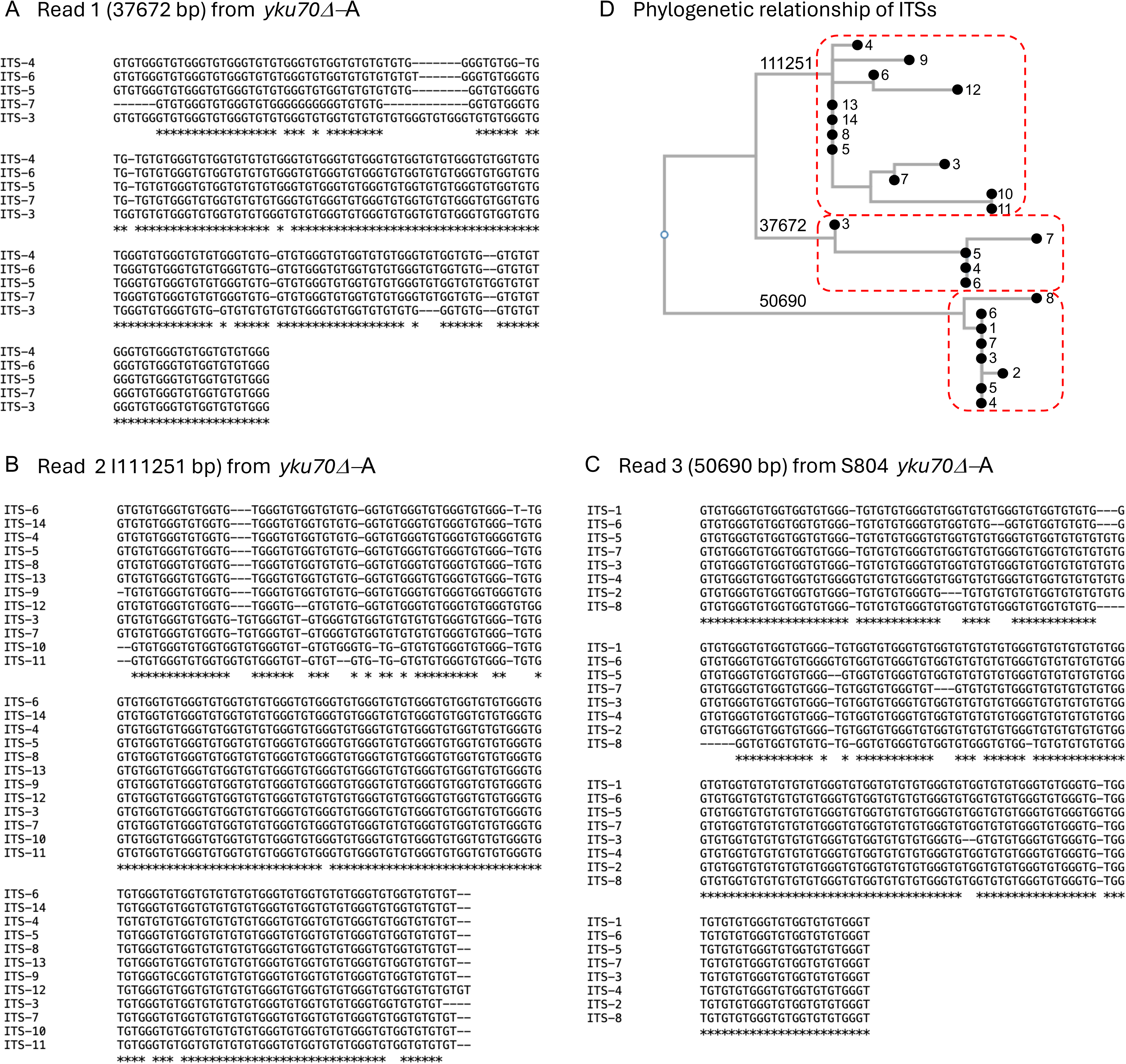
Multiple alignment of ITSs in tandem Y’s suggest that they are related by descent. A) The 52 bp ITS separating unique sequences from Y-1 of read 1 (37672 bp; Fig 5A) is identical in length and sequence to the parental ITS. ITSs 3-7 are similar in length and highly homologous in sequence. The poly-G stretch in ITS-7 is probably an artifact of Nanopore sequencing. B and C) ITSs from reads 2 and 3 (111251 bp and 50690 bp; Fig 5B and C) are highly similar within the cluster but differ between the two different clusters and D) differ from the ITS sequence of Read 1 (37672 bp; Fig 5D).

**S6 Figure:**
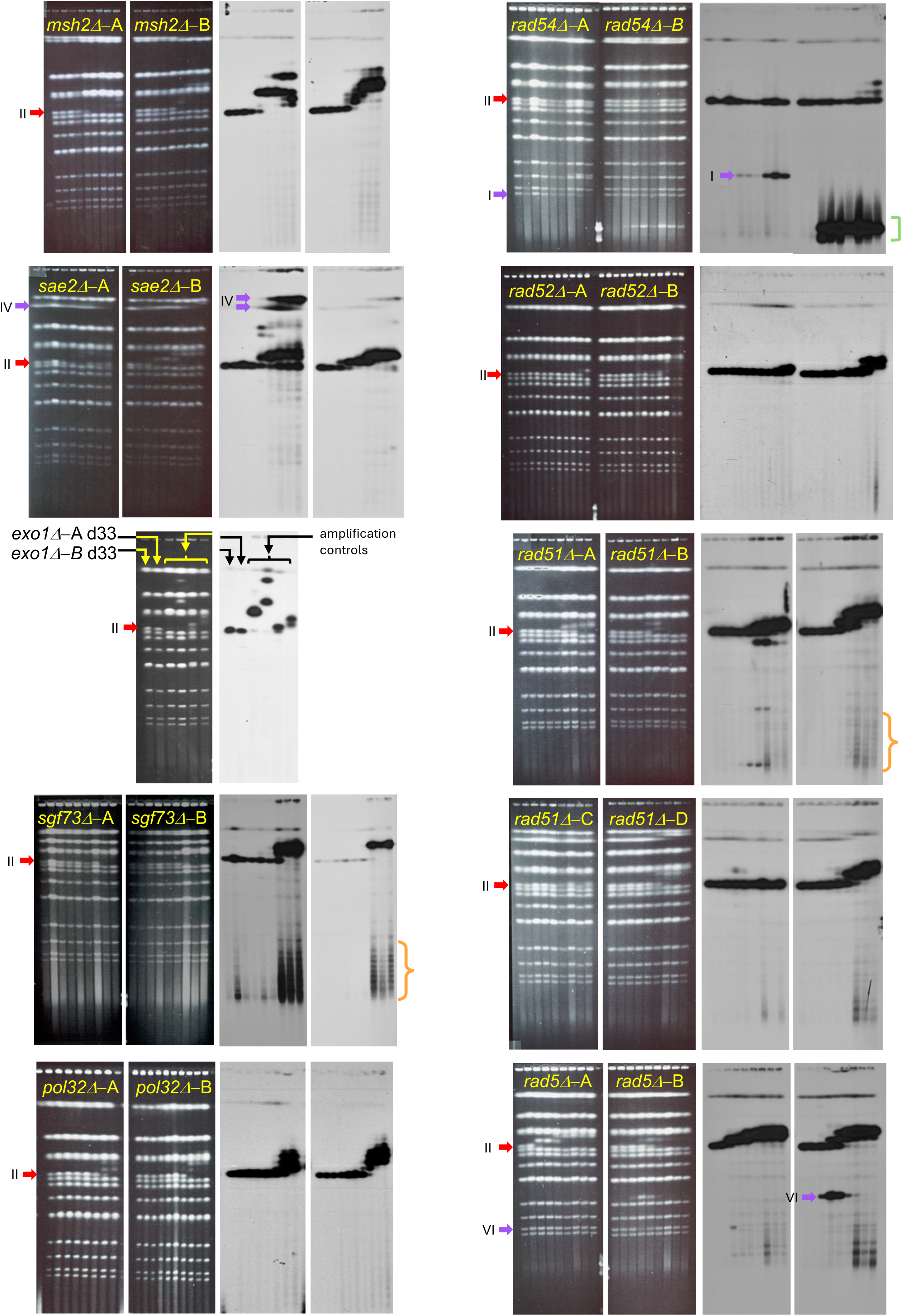
Characterization of chemostat populations of nine gene knockouts by CHEF gel electrophoresis and Southern blotting. Nine strains containing knock-out mutations in DNA maintenance genes were grown, independently in duplicate (or quadruplicate), in sulfate limiting chemostats for ∼200 generations. Cell populations were sampled at intervals and examined on CHEF gels (ethidium bromide image; left) followed by hybridization with *SUL1* (right). The two exceptions were the two *exo1ι1* chemostats, which were negative for *SUL1* amplification based on aCGH (see Fig 7). For these two isolates only the final day (day 33) was analyzed by CHEF gel to confirm the unaltered state of chromosome II. In each of the other cultures the major karyotypic change was an increase in the size of chromosome II (ethidium bromide-stained gel) resulting from amplification of a chromosomal fragment containing *SUL1* (Southern blots). In several population samples there is evidence that *SUL1* also moved to other chromosomes (I, IV and VI in *rad54ι1*–A, *sae2ι1*–A and *rad5Δ*-A, respectively; purple arrows) and of extrachromosomal supercoiled circular forms of *SUL1* amplicons (detected as topoisomers in *sgf73ι1*–A,B and *rad51ι1*–B, orange brackets) which are likely the product of pop-out recombination from tandem duplications. *rad54ι1*–B produced a linear extrachromosomal fragment (green bracket).

**S7 Figure:**
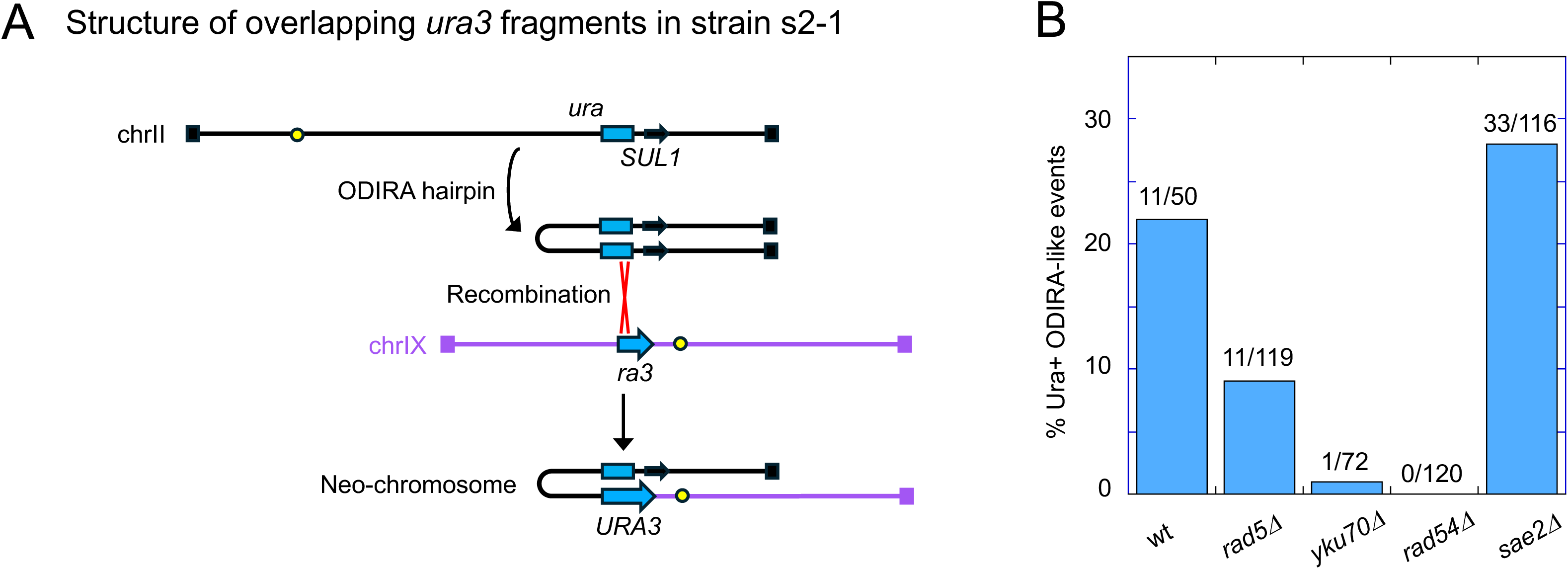
Quantification of ODIRA-like events using a split-*URA3* assay. A) Schematic diagram of chrII and chrIX in the strain s2-1. A 5’ fragment of the *URA3* gene (‘*ura’*) was inserted at the *SUL1* locus on chrII and an overlapping 3’ fragment (‘*ra3’*) was inserted to the left of the centromere on chrIX. The two major ways that cells recreate a functional *URA3* gene are by direct recombination between the two cassettes (not shown) or by the creation of an ODIRA hairpin from chrII-R and recombination of one of its two copies of *ura* with the *ra3* sequence on chromosome IX. This structure contains an inverted segment of chrII, upstream of the *SUL1* region, that is capped by the right telomeres of chrII and IX. B) Frequencies of independent ODIRA events were determined for each of four mutants (*rad5ι1*, *yku70ι1*, *rad54ι1*, and *sae2ι1*) and compared to the frequency found in wild type cells [2].

**S8 Figure:**
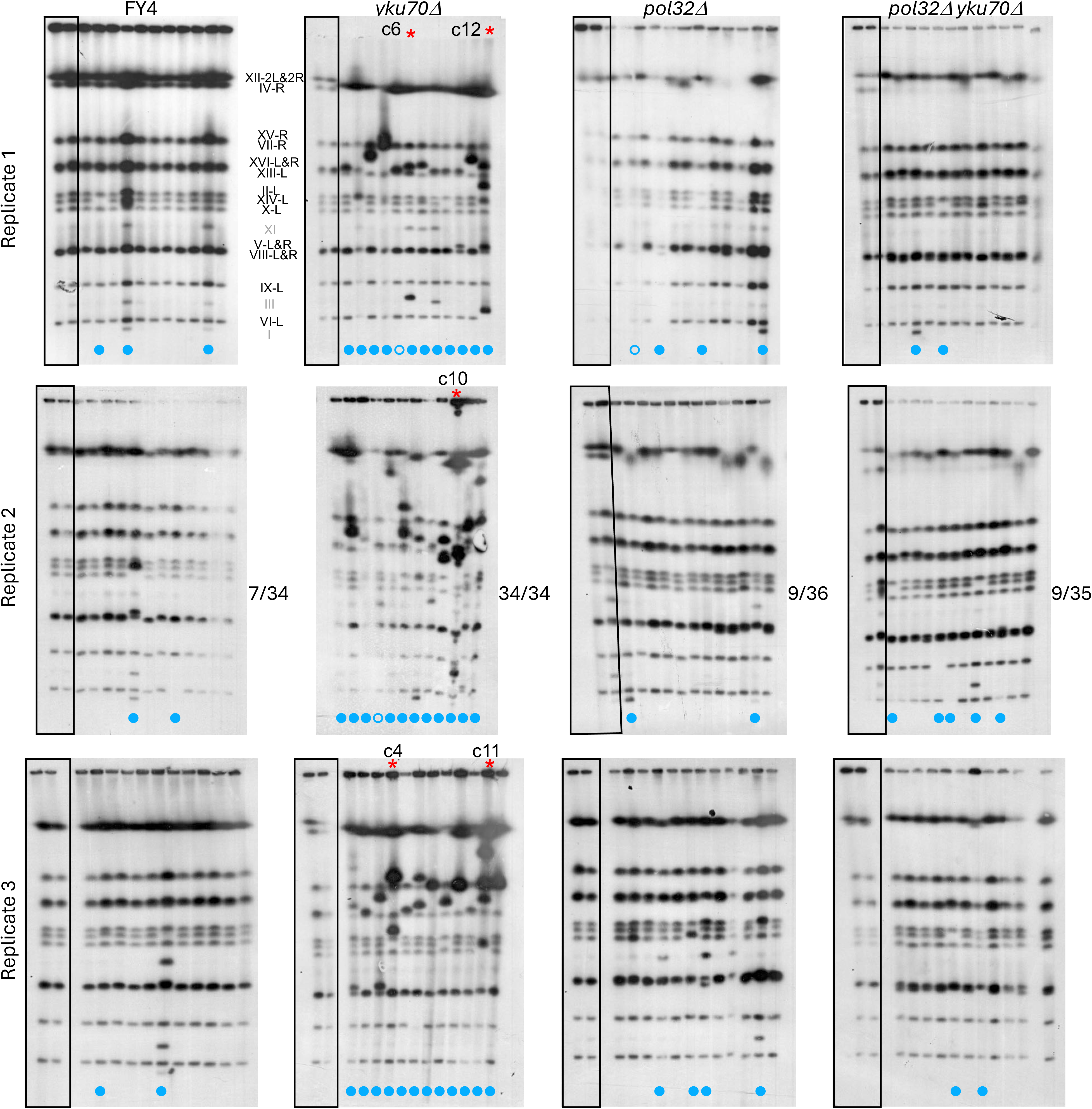
Technical replicates of serial transfer experiments with the wild type, *yku70ι1*, *pol32ι1,* and *yku70ι1 pol32ι1* strains. Three technical replicates of each 22-day serial transfer experiment were performed and 11-12 clones were examined from the last day of each culture. The Y’-probed Southern blots of the CHEF gels are shown. Results from replicate 3 for each of the four strains is repeated from Fig 8 for comparative purposes. Control lanes are boxed and clones with altered chromosomes are indicated by the cyan dots. (Hollow cyan dots indicate a second occurrence of a similar karyotypic change in that clone indicating it might not be a unique event.) Because the size of chromosome XII is dependent on the number of rDNA repeats and because Chr IV and ChrXII are not well resolved on these gels, we have not included potential changes to these chromosomes in our tally of clones with altered chromosomes. Clones analyzed by Nanopore sequencing are indicated by red asterisks and the results are shown in S9 Fig.

**S9 Figure:**
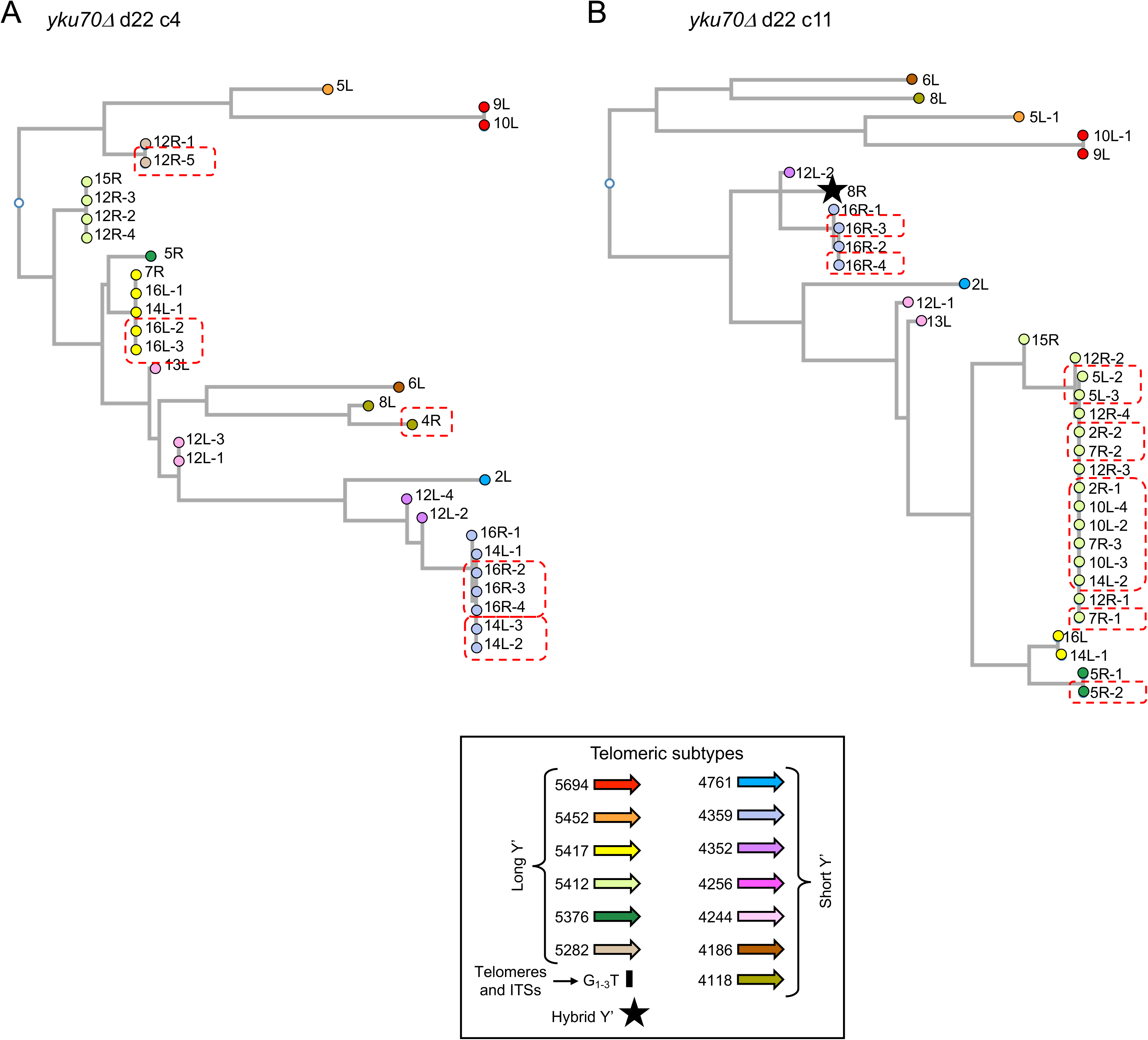
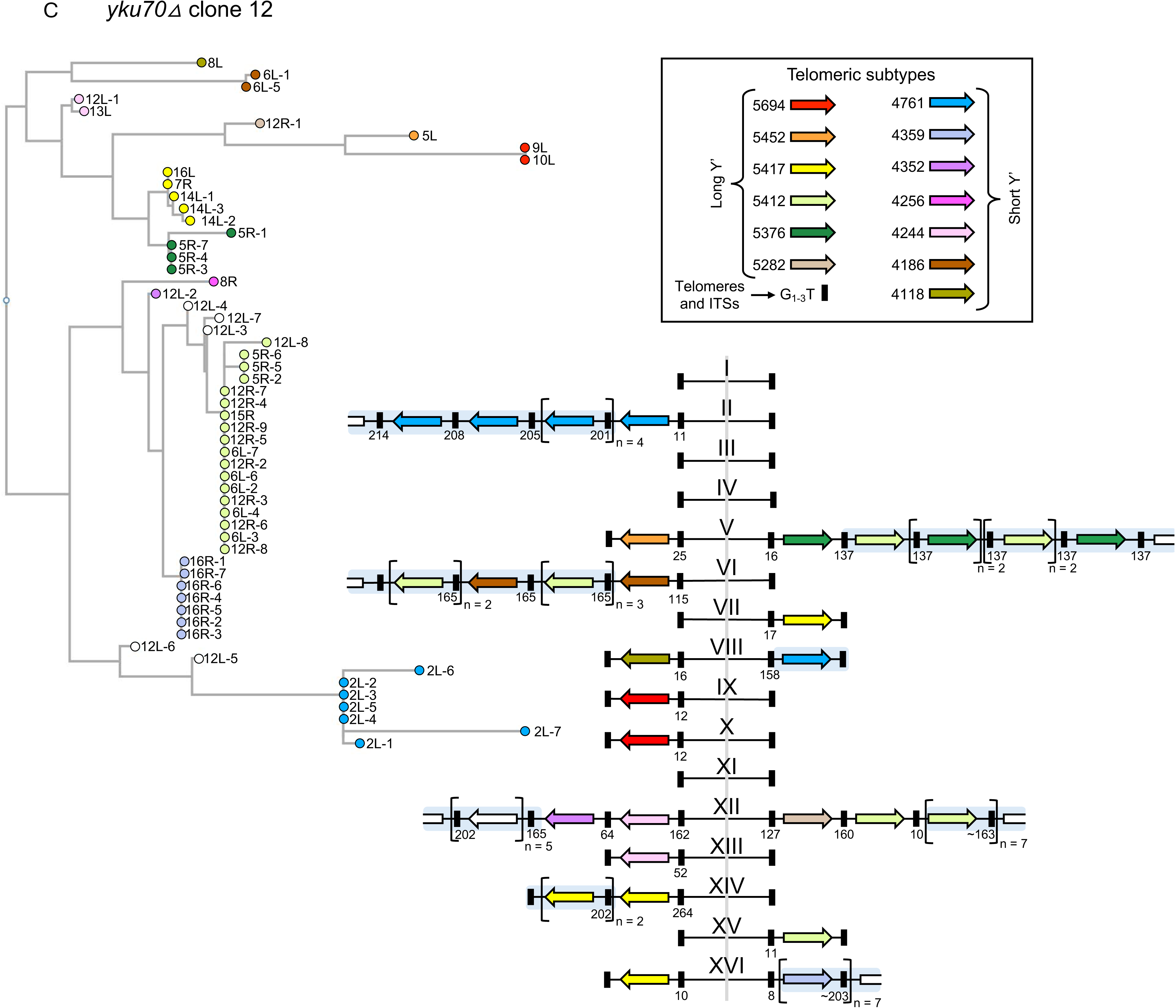
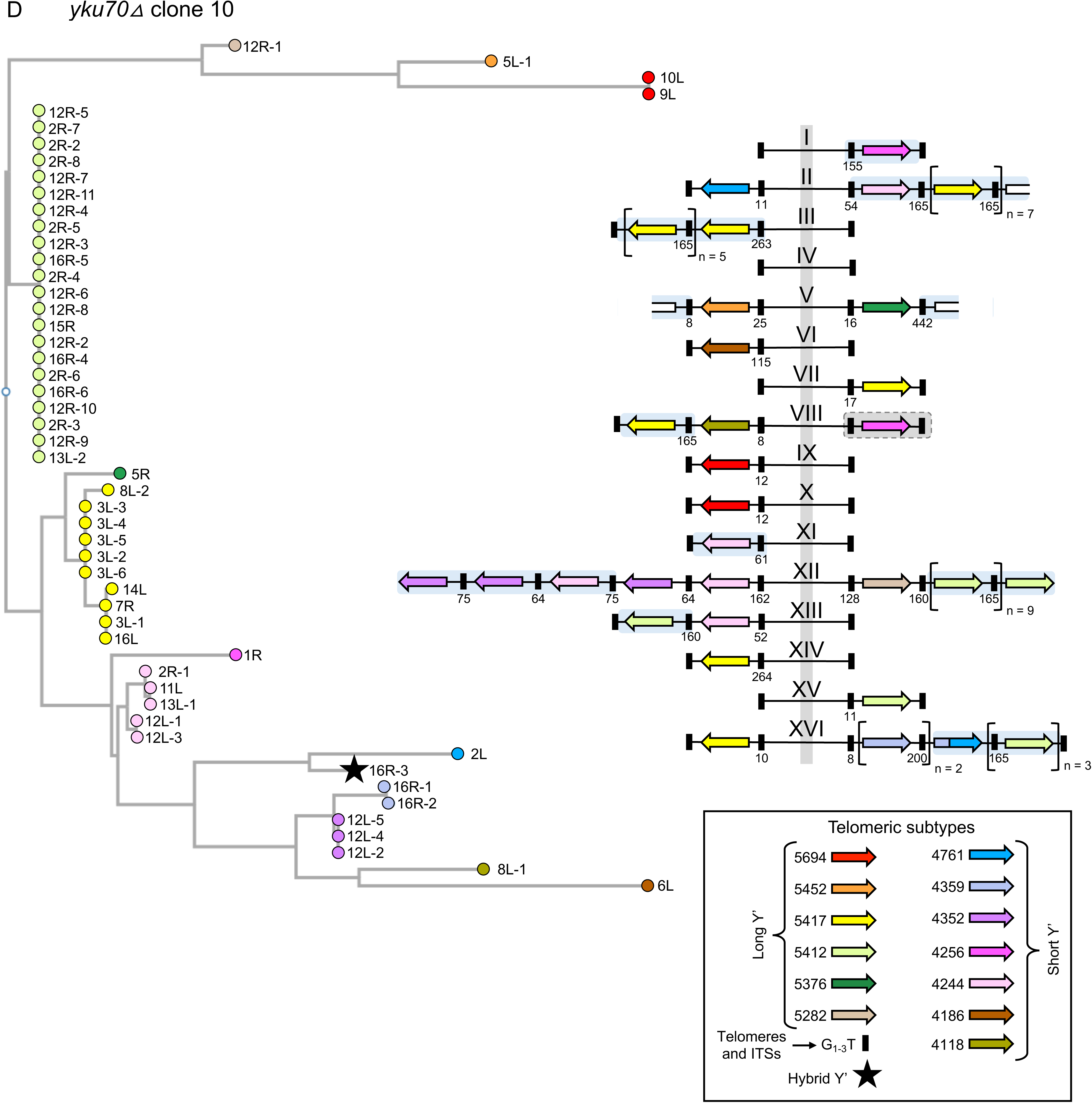
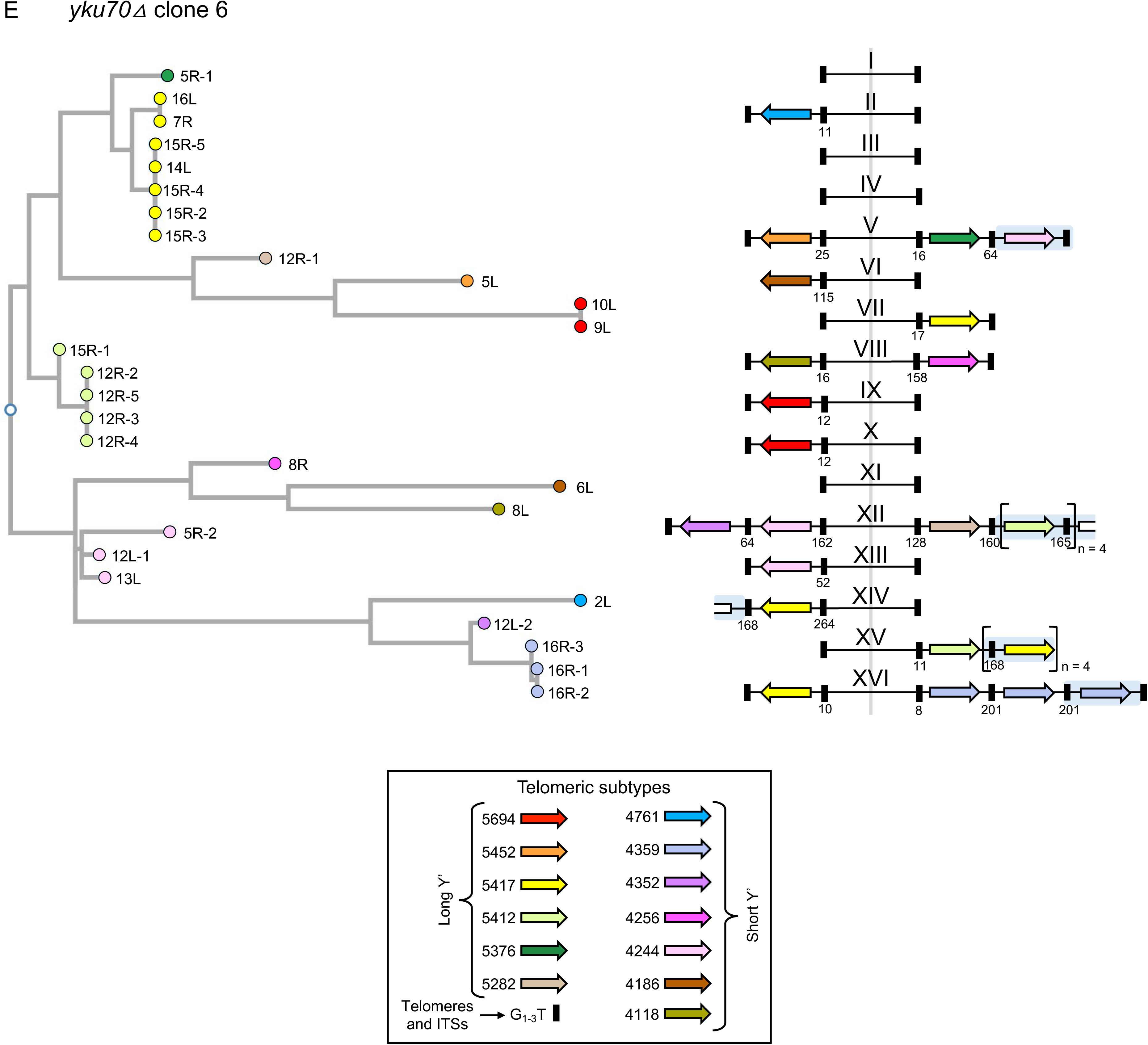
**Phylogenetic relationships of Y’s in clones derived from serial transfer experiments with *yku70****Δ* Clones 4, 11, 12, 10 and 6 (A-E, respectively) from day 22 of the serial passaging of the *yku70Δ* mutant (S8 Fig, red asterisks) were sequenced by Nanopore and telomeric amplification events were identified from individual reads and compared with the Y’s of the parental *yku70Δ* strain from day 0. The telomere maps were derived from the contigs with structural variants highlighted in blue shading. The origins of amplified Y’s were determined from the phylogenetic trees generated from CLUSTALW multiple alignments and are encircled by red dotted lines. See Fig 8 for telomere maps of clones c4 and c11. Two examples of hybrid Y’s were found: one on chrVIII-R in clone 11 (B; black star) and the second on chrXVI-R in clone 10 (D; black star).

**S10 Figure:**
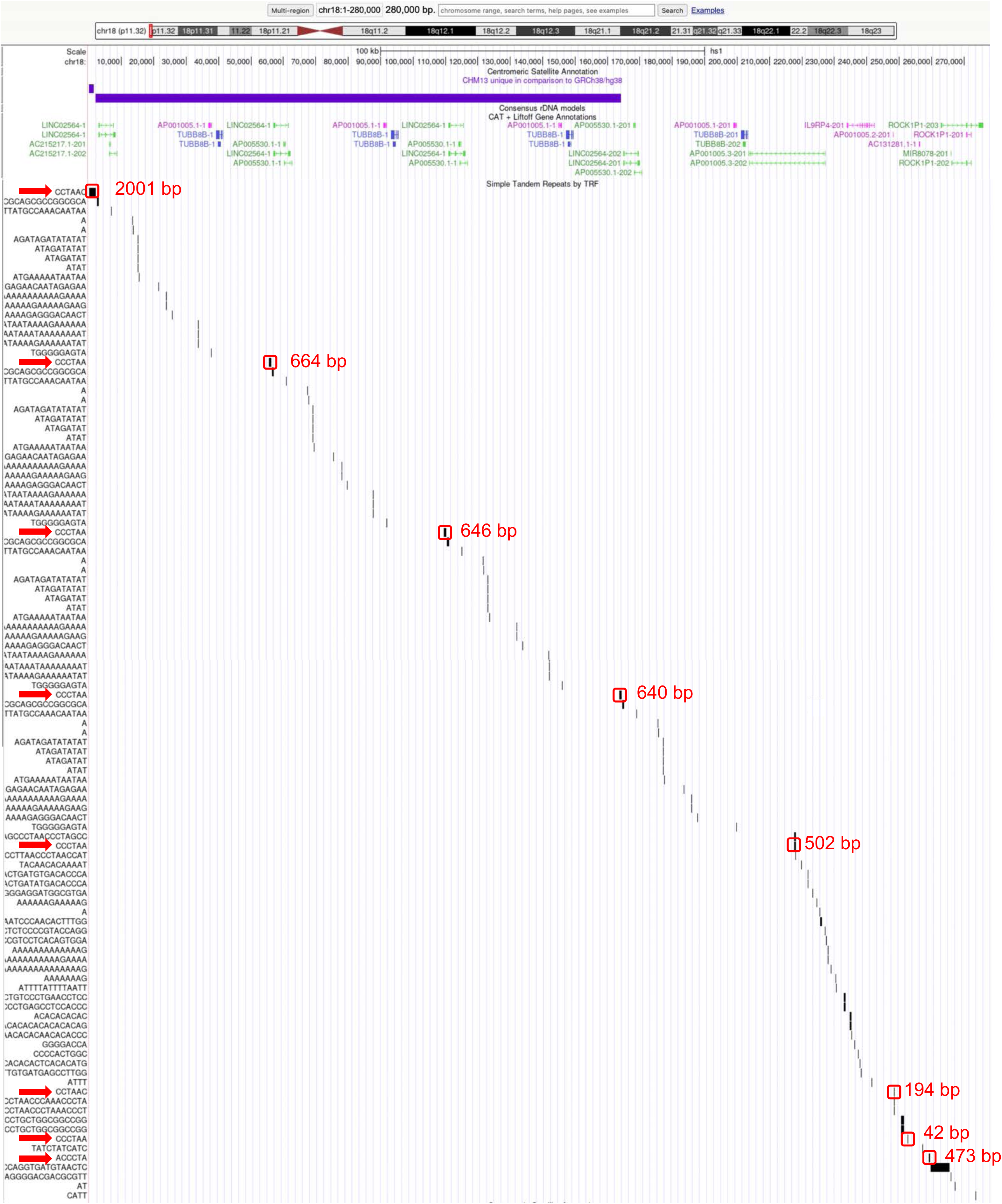
A screen shot from the T-to-T human genome browser showing the P-telomere of Chromosome 18. The ideogram of chromosome 18 is shown at the top with the P-telomere highlighted in red. The ∼275 kb telomeric region is expanded below. The horizontal purple line indicates the new sequences added to the human genome reference by the T-to-T project. Immediately below that are the positions of transcribed regions/known genes. The remainder of the image lists of all of the simple repeats found in this region of the genome and their relative positions. Included are the tracks of telomere repeats (CCCTAA, CCTAAC, CTAACC, etc.) which are labeled in red, with the lengths of those tracks in bp. The repetitive pattern generated by the other non-telomeric simple sequences highlights the near identity of the four 54 kb repeats.

---

**S1 Table.**
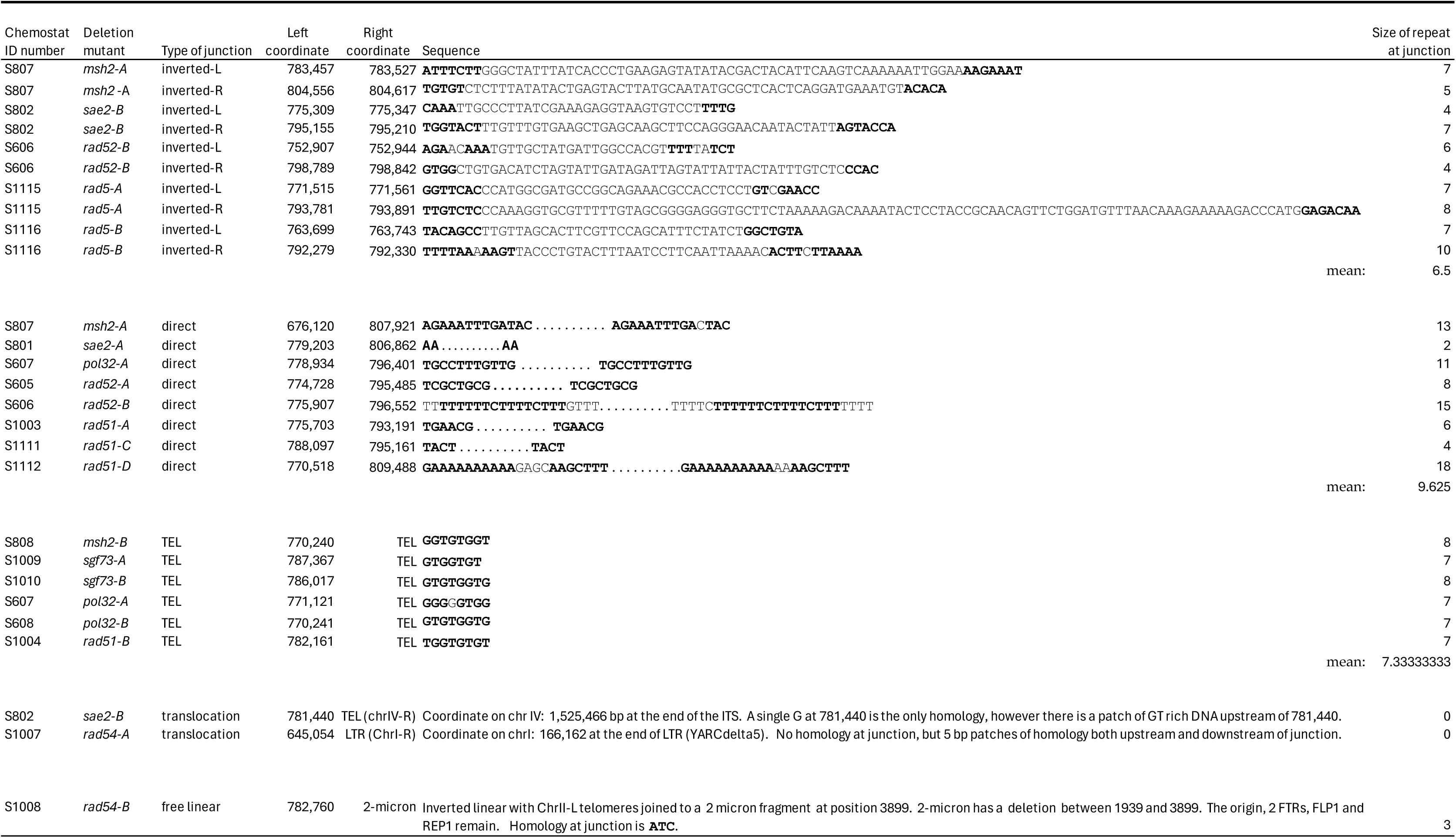
Junctions recovered from mutants in Fig. 3.

**S2 Table.** Summary of long-read sequencing runs.

| <u>Genotype</u> | <u>Alias</u> | <u>Total # of reads</u> | <u>Ave. read depth</u> |
| --- | --- | --- | --- |
| wt | FY4 | 259,655 | 126x |
| *yku70* $\Delta$ | *yku70* parent | 103,291 | 33x |
| *yku70* $\Delta$ | *yku70* Δ-A | 767,475 | 90x |
| *yku70* $\Delta$ | c10 | 253,646 | 113x |
| *yku70* $\Delta$ | c12 | 234,399 | 114x |
| *yku70* $\Delta$ | c4 | 112,130 | 49x |
| *yku70* $\Delta$ | c6 | 236,175 | 106x |
| *yku70* $\Delta$ | c11 | 337,611 | 122x |
| *ars228* $\Delta$ | clone 1475 | 117,143 | 91x |
| *msh2* $\Delta$ | S808 | 140,965 | 91x |
| *msh2* $\Delta$ | S807 | 228,384 | 139x |
| *pol32* $\Delta$ | S608 | 195,385 | 149x |
| *pol32* $\Delta$ | S607 | 242,580 | 141x |
| *rad51* $\Delta$ | S1112 | 235,662 | 65x |
| *rad51* $\Delta$ | S1111 | 786,141 | 204x |
| *rad51* $\Delta$ | S1004 | 280,123 | 150x |
| *rad51* $\Delta$ | S1003 | 289,533 | 132x |
| *rad52* $\Delta$ | S606 | 196,059 | 86x |
| *rad52* $\Delta$ | S605 | 174,117 | 85x |
| *rad54* $\Delta$ | S1008 | 159,449 | 78x |
| *rad54* $\Delta$ | S1007 | 104,682 | 54x |
| *rad5* $\Delta$ | S1116 | 158,311 | 86x |
| *rad5* $\Delta$ | S1115 | 159,438 | 97x |
| *sae2* $\Delta$ | S802 | 199,014 | 85x |
| *sae2* $\Delta$ | S801 | 1,246,025 | 149x |
| *sgf73* $\Delta$ | S1010 | 407,415 | 128x |
| *sgf73* $\Delta$ | S1009 | 97,093 | 28x |

